# Application of a new data management system to the study of the gut microbiome of children who are small for their gestational age

**DOI:** 10.1101/2024.06.11.598475

**Authors:** Felix Manske, Magdalena Durda-Masny, Norbert Grundmann, Jan Mazela, Monika Englert-Golon, Marta Szymankiewicz-Bręborowicz, Joanna Ciomborowska-Basheer, Izabela Makałowska, Anita Szwed, Wojciech Makałowski

**Author notes:** The authors contributed equally to the study (Felix Manske, Magdalena Durda-Masny). Author for correspondence: Wojciech Makałowski.

## Abstract

Microbiome studies aim to answer the following questions: which organisms are in the sample and what is their impact on the patient or the environment? To answer these questions, investigators have to perform comparative analyses on their classified sequences based on the collected metadata, such as treatment, condition of the patient, or the environment. The integrity of sequences, classifications, and metadata is paramount for the success of such studies. Still, the area of data management for the preliminary study results appears to be neglected.

Here, we present the development of a central data management system with an accessible web interface for the study of the gut microbiome in children who are small for their gestational age. We have called this system MetagenomicsDB. The web interface allows users, regardless of their bioinformatics expertise, to operate the system. The interface contains links to external resources in order to facilitate literature search, statistical analyses, and assessments of the potential pathogenicity of taxa. Preliminary study results are automatically quality-controlled and subsequently imported into a relational database. After exploration and optional filtering by the user, data are exported in a format directly suitable for follow-up analyses. Compared to a more conventional approach of storing the data in plain files, the automated quality control and database storage offered by MetagenomicsDB provides an enhanced quality assurance of the produced study results. This is especially true in a collaborative setting. Also, the automation of the data transfer from the format of the input data to the format needed for downstream analyses makes basic statistical and bioinformatic analyses accessible. The web interface not only allows us to perform our internal analyses, but will also facilitate transparent sharing of the complete study results at publication time with reviewers and the general public. We demonstrated the viability of this approach by making a subset of our preliminary study results already publicly accessible: https://www.bioinformatics.uni-muenster.de/tools/metagenomicsDB

In the context of our study, our system provided more flexibility to conduct study-specific analyses and to integrate specific external resources than existing and necessarily more generic solutions. Our system is not yet ready to be widely applicable out-of-the-box for any microbiome study. However, we expect that due to the modular concept, the tests, and the extensive description of the underlying rationale, large parts of our implementation can be adapted for future projects in a fraction of the time needed to develop a new data management system from scratch. Thus, we report our endeavors in order to motivate the application of data management systems at the scale of single studies in microbiome research.

## 1 Introduction

Traditionally, the human gut microbiome was analyzed using culture-based approaches (Aries et al. 1969; Finegold, Attebery, and Sutter 1974; Moore and Holdeman 1974). Technological advances enabled the identification of bacterial communities by whole-genome sequencing (metagenomics) or the analysis of marker genes (Escobar-Zepeda, Vera-Ponce de León, and Sanchez-Flores 2015), which provided a new understanding of microbial habitats: 80% of bacterial operational taxonomic units (OTUs) in the human gut, which were detected by analysis of the 16S rRNA gene, had not been cultured before (Eckburg et al. 2005). Despite new and improved efforts towards culture-based approaches (Lagier et al. 2012), sequenced-based analyses are still commonly used to investigate diseases (Fujimura et al. 2016; Matson et al. 2018), the impact of external factors such as diet (David et al. 2014; Bäckhed et al. 2015), and the mode of delivery on the microbiome (Dominguez-Bello et al. 2010; Bäckhed et al. 2015).

In an ongoing prospective study, we are investigating the role of the gut microbiome in determining birth weight and weight gain for term children who are small for gestational age (SGA) throughout the first year of life by using full-length 16S rRNA gene sequences. To the best of our knowledge, a similar study has not been conducted so far. Low birth weight has been shown to increase the risk of developing various adverse conditions later in life, such as cardiometabolic disorders (Barker et al. 1993; Ravn Knop et al. 2018), attention-deficit/hyperactivity disorder (Beer et al. 2022), and learning difficulties (O’Keeffe et al. 2003). In this study, we define SGA children as low-birth-weight children with a weight-for-age z-score (WHO Multicentre Growth Reference Study Group 2006, 302–3) of less than negative two at birth (World Health Organization 2022, 1216). In 2022, approximately 0.3 million live births occurred in Poland (3.9 million in the European Union) (https://doi.org/10.2908/TPS00204; accessed 2024/04/17). Given that a z-score of -2 is roughly equal to the 2.3rd percentile, we expect that almost 7,000 children in Poland (approximately 90,000 children in the European Union) have been born with SGA in 2022.

During our study, we noticed that a major challenge of microbiome studies for laboratory scientists is the data analysis. In our study, we had produced patient and sample metadata, sequencing results, and classifications of the sequences which needed to be stored in an accessible and secure way. Also, our analyses required the use of weight-for-age growth standards from the World Health Organization (WHO). The critical aspect of storing the different data types was not only to ensure the consistency of each, but also to maintain the correct connections between them. Subsequently, statistics needed to be performed on samples or patients of interest. This required us to subset the stored data and to provide them in a specific format to the statistical software. All of these operations had to be performed by laboratory scientists. Thus, the use of command line interfaces or the manual execution of a collection of in-house scripts was not an option.

In order to cope with the aforementioned challenges, we designed a central data management system with a web-based graphical interface, the MetagenomicsDB. It features a fully automatic import and quality control of patient and sample metadata from spreadsheets produced by popular programs, such as Excel (https://www.microsoft.com/en-us/microsoft-365/excel; accessed 2024/02/29) or LibreOffice Calc (https://www.libreoffice.org/; accessed 2024/02/29), but also from tab-delimited text files. This enhances use for collaborative projects with medical practitioners, lab scientists, and bioinformaticians, who typically prefer different input formats. Sequencing results in the FASTQ format and taxonomic classifications from MetaG (Manske, Grundmann, and Makałowski 2020) software are retrieved from a common directory and automatically connected to the respective samples in the database using unique identifiers from the spreadsheet. With the graphical interface, investigators can export data of interest from the relational database to MicrobiomeAnalyst (Lu et al. 2023) for statistical analyses.

We designed the core components of MetagenomicsDB to be as flexible as possible in order to increase the value of our system for future studies. Components were built using a modular concept and were supplemented with tests. This permits the adaptation of existing functionalities or the addition of new ones by modifying existing or adding new components. In line with this, we used a relational database management system to keep the user interface inert to alterations of the physical and logical structure of the stored data (Codd 1985, ID/8). We reported our endeavors by creating a central data management system by the example of our SGA study in order to motivate the application of databases as central components for managing data produced by ongoing studies. We have made MetagenomicsDB publicly accessible with a reduced subset of our study data: https://www.bioinformatics.uni-muenster.de/tools/metagenomicsDB.

## 2 Data produced by the exemplary SGA study

Our ongoing prospective study investigated the potential influence of the gut microbiome on birth weight and weight gain in SGA children. Only single birth children who were born on time and healthy (apart from the condition under study) were eligible (Supplementary Methods 1). 65 children of Polish descent were regularly examined at a maximum of 8 time points within the first year of life (Supplementary Methods 1). At birth (“meconium” sample), we recorded factors related to the pregnancy and neonatal development, such as potential illnesses of the mother and her weight gain during pregnancy (Supplementary Table S1; Supplementary Methods 2). At the sampling time points, patients were weighted, stool samples were collected, and confounding factors to the gut microbiome (such as antibiotics use, probiotics consumption, and feeding habits of the patients) were recorded (Supplementary Table S1; Supplementary Methods 2 and 3). The measurements and the patient and maternal metadata were stored in an Excel spreadsheet (Supplementary Table S1). Importantly, spreadsheet provided a key which connected the “pass” reads obtained from nanopore sequencing of the stool samples (Supplementary Methods 3.3) to the respective entry in the spreadsheet. This key consisted of the run and barcode numbers of the full-length 16S rRNA gene sequences (column: “number of run and barcode”, Supplementary Table S1). Each sequencing run additionally contained an internal control to verify that library preparation and sequencing did not introduce contamination (Supplementary Methods 3.3). The control was always assigned to the special barcode 99 of the respective run (Supplementary Methods 5.3.1).

All “pass” reads (Supplementary Methods 3.3) were initially filtered for residual human contamination using the T2T-CHM13v2.0 (RefSeq (Sayers et al. 2024) accession: GCF_009914755.1) human reference genome in order to improve the accuracy of the taxonomic classifications and help protect the privacy of patients (Supplementary Methods 4.1). The remainder was classified by MetaG using RDP (Cole et al. 2014) version 11.5 (Supplementary Methods 4.2). The program and database name of the classification workflow were added to the spreadsheet (columns: “program” and “database”, Supplementary Table S1). Both FASTQ files and their classifications were stored in a common directory with subdirectories named after the respective key in the spreadsheet (Supplementary Methods 5.3.1; column: “number of run and barcode”, Supplementary Table S1). In order to evaluate the growth of the children by their weight-for-age z-scores, we downloaded the respective growth standards for boys (https://cdn.who.int/media/docs/ default-source/child-growth/child-growth-standards/indicators/weight-for-age/expanded- tables/wfa-boys-zscore-expanded-tables.xlsx; accessed 2024/01/30) and for girls (https://cdn.who.int/media/docs/default-source/child-growth/child-growth-standards/ indicators/weight-for-age/expanded-tables/wfa-girls-zscore-expanded-tables.xlsx; accessed 2024/01/30) from the WHO.

## 3 MetagenomicsDB: a central data management system

We designed a central data management system in order to organize, store, and analyze our preliminary study results. We called this system MetagenomicsDB. It consists of three main components (Figure 1): an import unit which checks the integrity of the input data and synchronizes the quality-checked data with the database (Section 3.1). Preliminary study results and associated metadata were stored using the relational database management system PostgreSQL (https://www.postgresql.org/; accessed 2024/03/17) (Section 3.2). Another unit exports a user selection of the database contents in a format suitable for downstream statistical analyses in MicrobiomeAnalyst (Section 3.3). All components can be operated using a web-based graphical interface (Figure 1), which allows users, regardless of their bioinformatics expertise, to store, explore, and export the preliminary study results (Section 3.4).

**Figure 1:**
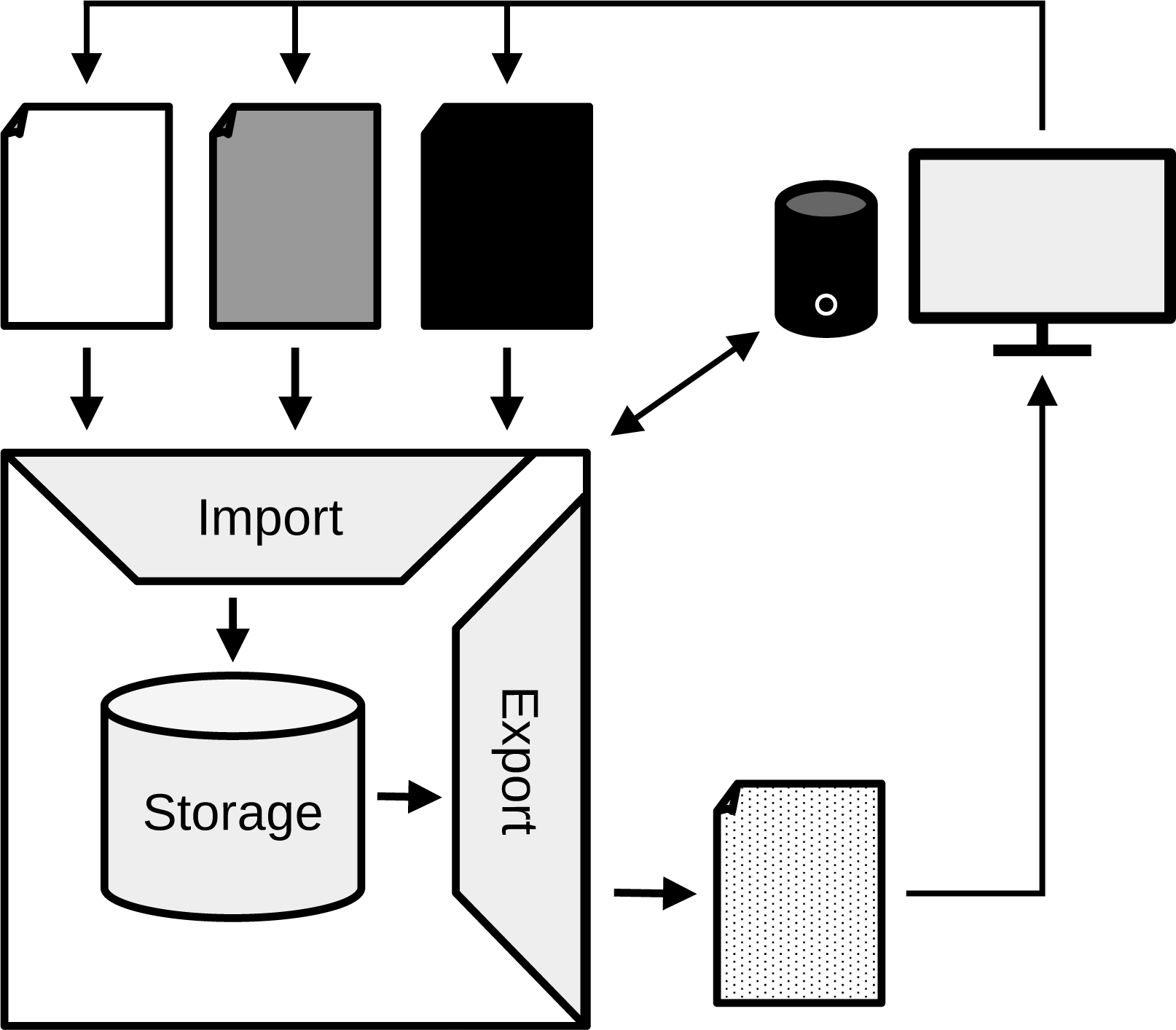
Components of MetagenomicsDB. Different inputs, illustrated by the three differently colored files, are quality-checked and synchronized with the storage unit with the help of a general import unit. A user selected subset of data is then exported by the respective component in a format suitable for downstream analyses (dotted file). MetagenomicsDB is operated using a web-based graphical interface (computer terminal), which enables users to upload input data for the synchronization with the database, explore the data contents, and export the results. This figure was generated with LibreOffice Draw 7.0.4.2 (https://www.libreoffice.org/discover/draw/; accessed 2024/03/22).

For future microbiome studies, we aimed to design MetagenomicsDB as a non-disposable data management system (Supplementary Methods 5.4). Thus, the import and export units (Figure 1) were built modular and supplemented with tests. This will allow us to reuse universal functions which we expected to be useful across projects, for example functions parsing FASTQ files or spreadsheets. Other more high-level functions, such as definitions of the columns which were supposed to be extracted from the spreadsheets, necessarily had to be project-specific. Due to the modular concept, project-specific functions can be easily replaced with their successors for future projects, considerably reducing the workload for follow-up projects. Modularity also facilitates the extension of the current functionalities to fit changed demands. For instance, only classifications from the MetaG classifier were processed in this study (Supplementary Methods 4.2). In future projects, the use of other popular classifiers, such as QIIME 2 (Bolyen et al. 2019) or Kraken 2 (Wood, Lu, and Langmead 2019), could be of interest. To achieve support for more classifiers, the only major change to the import unit of MetagenomicsDB is the addition of a new module. The module parses the specific output file format of the new classifier and converts it to the common internal data representation of our pipeline. In line with the modular concept, we designed a flexible database schema (Section 3.2) and used the relational database management system PostgreSQL as the storage component, since relational database management systems keep the user interface inert to alterations of the physical and logical structure of the stored data (Codd 1985, ID/8). In the following sections, the three components of MetagenomicsDB (Figure 1) and its web-based graphical interface will be presented. An exhaustive description of the implementation and the underlying rational is presented in Supplementary Methods section.

### 3.1 Import of the study data

A single program called importSGA.pl (commit: 68b05c6; https://github.com/IOB-Muenster/MetagenomicsDB/blob/main/bin/importSGA.pl; accessed: 2024/06/07) was responsible for the import and quality control of the Excel spreadsheet with the measurements and metadata, the FASTQs, the MetaG classifications of reads (“calc.LIN.txt” files), and the WHO weight-for-age growth standards (Section 2; Supplementary Table S6). The main duties of this program were reading and, if necessary, extracting the input files, parsing the contents, performing quality control to reduce human error, and synchronizing the data with the database. In the following, we will briefly present the general algorithm of our program for the handling of different input file types. For the sake of simplicity, we will use the term “measurements” for all contents of the Excel spreadsheet.

The WHO standards were only inserted once at the beginning of the project and read by our program as is (Supplementary Methods 5.3.1). We processed the classifications from the MetaG software provided in the “calc.LIN.txt” files (Supplementary Methods 5.3.1). The files contained the classifications per read and per rank (domain, phylum, class, subclass, order, suborder, family, genus, species, and strain). For each of these ranks, reads that could not be classified at all and reads that could not be classified from the respective rank onward were assigned to the special taxon “UNMATCHED.” Reads were assigned to the taxon “FILTERED” over all ranks if the read appeared in the FASTQ files but not in the “calc.LIN.txt” files. Since we had removed reads with human contamination prior to the taxonomic classification (Section 2), the “FILTERED” taxon represented reads identified as human contamination. The taxon name was null in the case no name was available (Supplementary Methods 5.3.1). After an optional extraction, our program concatenated all FASTQ files (Supplementary Methods 5.3.1) found in a subdirectory. However, we enforced that files had to have distinct names (not considering the file extension) in order to avoid concatenating duplicate data. By comparing the read IDs in both files, importSGA.pl verified that FASTQ files and the respective classification files were matching. Sequences, quality strings, read IDs, and metadata from the headers were extracted in order to store them in the database (Supplementary Methods 5.3.1).

Only certain columns were extracted from the Excel spreadsheet containing the measurements (Supplementary Table S7). The vast majority of ignored columns represented measurements whose values could be calculated based on values of other measurements. We named these derived measurements (Supplementary Table S4). Derived measurements were calculated in the database itself (Section 3.2; Supplementary Methods 5.1.2) in order to avoid update anomalies. For example, the weight-for-age z-score of a patient depended on his or her birthdate, the patient’s sex, the sample collection date, the patient’s body mass, and the WHO growth standards (Supplementary Methods 5.1.2). If the z-score was only read from the input, a later update of the patient’s birthdate in the database would leave the database in an inconsistent state, as the dependent z-score would not have been automatically updated. Some columns in the spreadsheet contained values which were encoded by numbers. importSGA.pl automatically translated these numbers into the referenced values (Supplementary Methods 5.3.2).

The spreadsheet connected the sequences and classifications to the respective samples by providing a unique key in the form of the run and barcode of the sequences (Section 2). These keys only referred to the genuine stool samples and not to the internal controls from each run. In order to link the control to all samples of the respective sequencing run, we internally duplicated each sample. One copy was labeled as control and only connected to the sequences from the control barcode 99. The other copy was labeled as the case sample and was assigned to the genuine stool sample sequences and the extracted measurements from the spreadsheet (Supplementary Methods 5.3.2).

The import program connected to the database with the service name “metagdb” provided in the PostgreSQL configuration file “.pg_service.conf” on the host machine (Supplementary Methods 5.3.2), which made the application easily portable. We ensured that only one instance of our program was running at the same time by attempting to acquire an exclusive transaction-level advisory lock. If the lock was already held, the new instance of the program was terminated (Supplementary Methods 5.3.2). All database manipulations were performed in the same transaction that was used to acquire the lock, thus all changes could be completely rolled back in the event of an error. Inserts into and updates of the database were conducted in batches for improved performance over the network (data not shown).

After connecting to the database, our script filtered duplicates in the input data and autonomously decided how to synchronize the input files with the database by using the unique constraints of the relations (Supplementary Methods 5.1.1 and 5.3.2). This allowed us to simultaneously provide old records as well as records that were supposed to be inserted or updated as input data. Generally, however, it was advantageous to limit input files to new or changed records in order to reduce the calculation time. Nevertheless, even in the case of such a minimal synchronization, old data were mandatory when they were needed to establish the correct connections between the data types (Supplementary Methods 5.3.2). For example, it was not possible to insert FASTQ files by themselves, as the measurement spreadsheet was essential to establish the connection between the sequences and the respective sample. Similar considerations were applied to the classification files which depended on the sequence files and thus also on the measurement spreadsheet (Supplementary Methods 5.3.1 and 5.3.2).

Overall, initializing the database from scratch with the WHO growth standards, 65 patients, 986 samples, and more than 3.8 million sequences (of which over 97% remained after filtering for human contamination), required roughly 3 hours on a Debian 11 server (2 AMD EPYC 7452 CPUs, 1 TB RAM) with access to the input data and the database over the network. The most time-consuming step was the parsing of FASTQs and the subsequent insertion of the sequence data, which accounted for more than half of the total calculation time.

### 3.2 Data storage

Data were stored in a PostgreSQL 14beta2 database. The schema consisted of ten relations (Figure 2) and five materialized views. The database schema is presented in depth in the Supplementary Methods (Supplementary Methods 5.1.1 and 5.1.2). In this section, we will focus on the general properties of the database schema. Names of views and relations will be printed in italics to distinguish them from regular English words.

**Figure 2:**
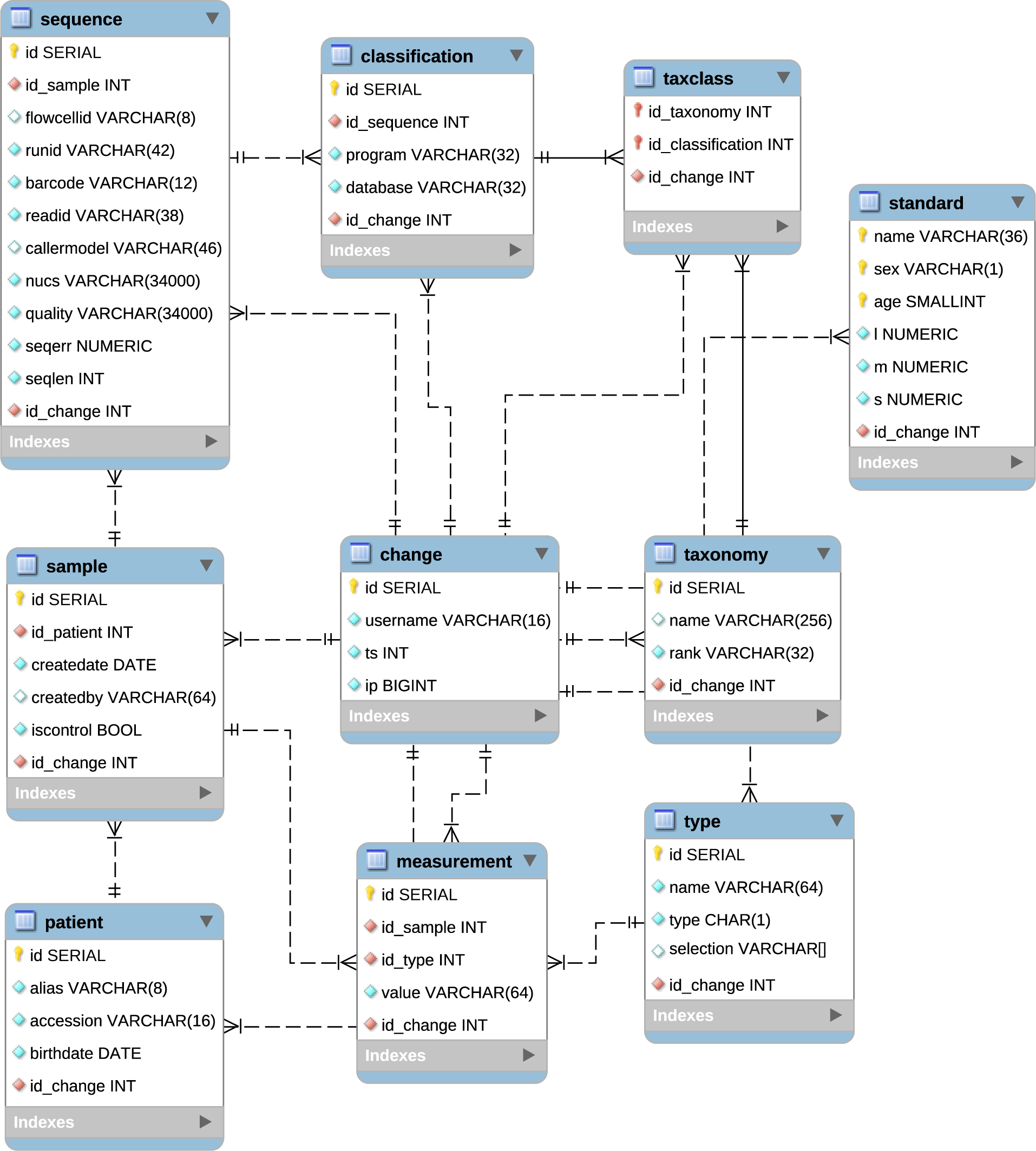
Relations in MetagenomicsDB. The key symbol indicates a primary key and the diamond symbolizes a regular column. Foreign keys are depicted by red symbols and turquoise shading indicates “not NULL” constraints. In our schema, all foreign keys also had a “not NULL” constraint (commit: 68b05c6; https://github.com/IOB-Muenster/ MetagenomicsDB/blob/main/www-intern/db/schema.sql; accessed 2024/06/07). Connections between the relations are shown by lines (dashed for non-identifying relationships). This database schema was visualized with MySQL Workbench 6.3.8 build 1228 CE Community (https://dev.mysql.com/downloads/workbench/; accessed 2024/03/15) and edited with Inkscape 1.0.2 (https://inkscape.org/; accessed 2024/03/15).

In our study, each patient in the *patient* relation was sampled at up to eight time points. Thus, individual patients were assigned to a maximum of eight genuine samples and eight control samples in the *sample* relation (Sections 2 and 3.1). The controls represented the results from sequencing the internal control sample of each run (Supplementary Methods 3.3) and were identified by the value in the column “isControl” in *sample* (Figure 2). Controls had no measurements, as opposed to the genuine samples. In order to keep the database schema flexible with regards to the number of stored measurements, we stored single measurements as rows in the *measurement* relation (Figure 2). This approach supported an arbitrary number of measurements per sample. The values of static measurements were only recorded once and never changed. These were values related to the mother (e.g.: maternal body mass before pregnancy), the pregnancy (e.g.: pregnancy order), and the birth (e.g. birth mode). The sex of the patient was also considered static (Supplementary Table S3). In order to reduce redundancy, we assigned static measurements only to the first sample (“meconium” sample) from each patient. Variable measurements, such as the patient’s body mass, were recorded during each of the appointments and were assigned to the respective samples (Supplementary Table S3). Derived measurements were only calculated in the views (see below). Measurement names were normalized by the *type* relation. The relation additionally contained the expected input type of the measurement value and, if appropriate, an array of possible values (“type” and “selection”, Figure 2). This information was used in an internal administration web interface in order to check the input values and to generate drop-down menus. Due to the storage of single measurements as rows in *measurement*, it was not possible to enforce specific constraints on the data type of values at the level of the database (Figure 2).

Each sample could be linked to multiple sequences. In the *sequence* relation, we calculate the read length and average sequence error of each entry (“seqlen” and “seqerr”, Figure 2) using stored generated columns. These helped to reduce redundancy, as read length and average sequence error could be directly inferred from the sequence and the quality string of each read, respectively (Supplementary Methods 5.1.1). Both values were later used by the views to calculate maximum, minimum, and average values for the sequence length and the average read quality of sequences supporting a specific taxon (Supplementary Methods 5.1.2). Each read could be classified with different programs and databases. We recorded this information in the *classification* relation (Figure 2). In our exemplary study, however, all reads were classified by MetaG using the RDP database (Supplementary Methods 4.2). The taxa from the classifications were stored separately by name and rank in the *taxonomy* relation, which was linked by the associative table *taxclass* to *classification* (Figure 2). The taxa in *taxonomy* were unique considering the combinations of name and rank (Supplementary Methods 5.1.1). However, in the course of the study, we found that RDP contained taxa with the same name and rank that came from different lineages (data not shown). Thus, even though taxa were considered identical by the database, they could sometimes represented distinct biological entities. We accounted for this fact in the web interface (Section 3.4). The *change* relation recorded information about the submitter of each record, which we used internally to track changes (Figure 2). Isolated from the rest but *change* was the *standard* relation (Figure 2), which contained information from the WHO growth standards (Supplementary Methods 5.3.1) and was needed to calculate the weight-for-age z-scores in the views (see below).

We used materialized views to calculate the derived measurements (such as the aforementioned weight-for-age z-score in *v_measurements*), to aggregate data for the export to MicrobiomeAnalyst (*v_lineages*, *v_samples*, and *v_metadata*), and for the functionalities of the web interface (Section 3.4; Supplementary Methods 5.1.2). Materialization was chosen in order to improve the speed of queries and views were automatically refreshed when necessary by importSGA.pl. Apart from the derived measurements (Supplementary Methods 5.1.2), the views also calculated a time point string, which expressed the sample collection date relative to the birthdate of the respective patient (*v_samples*). This allowed for convenient comparisons of samples from different patients at the same time point after birth. The definitions of time points were fuzzy in order to account for minor variation in the sample collection date (Supplementary Methods 5.1.2). For example, samples taken between 354 and 376 days were labeled as the 1 year (“1y”) sample. This increased flexibility came at the expense of not being able to enforce uniqueness of sampling time points at the level of our database. Additionally, it was not guaranteed that a time point string was assigned to each sample collection date, as time points were discrete. However, importSGA.pl enforced these two constraints when inserting or updating samples (Supplementary Methods 5.1.2). Other important information that was calculated in the views was the read count per sample (*v_samples*) and the assembly of the taxonomic classifications for a read into a complete lineage (*v_lineages*). Furthermore, minimum, maximum, and average values were calculated for the sequence length and the average read quality of sequences supporting a taxon identified by a specific program and database (*v_taxa*) (Supplementary Methods 5.1.2).

### 3.3 Export for statistical analyses

MetagenomicsDB provided an export of data to the “Marker Data Profiling” workflow (https://www.microbiomeanalyst.ca/MicrobiomeAnalyst/upload/OtuUploadView.xhtml; accessed 2024/02/01) of MicrobiomeAnalyst in order to facilitate downstream statistical analyses. MicrobiomeAnalyst is a tool that can use (among others) taxonomic classifications together with sample metadata to perform statistical analyses and functional predictions in a graphical web interface (Lu et al. 2023). When provided database IDs for samples of interest, exportSGA.pl (commit: 68b05c6; https://github.com/IOB-Muenster/MetagenomicsDB/blob/main/bin/exportSGA.pl; accessed: 2024/06/07) exported classifications and sample metadata to a ZIP archive (Supplementary Methods 5.2; Supplementary Table S5). By default, we excluded the “FILTERED” taxon and skipped any sample associated with the internal controls (Supplementary Methods 5.2). We only exported samples that had classifications and wrote the number of samples which were not exported to a log file in the archive (“microbiomeanalyst_WARNINGS.txt”). The raw counts of supporting reads for each lineage across all exported samples were stored in “microbiomeanalyst_otu.txt”. Lineages and samples were identified by unique keys. The keys for lineages contained the term “OTU” followed by a counter. The counter was based on the sorted order of lineages in the respective export. Thus, the counter was stable between multiple exports of the same samples (assuming the database contents remained unchanged), but was not lineage-specific across exports with different samples. Each ID was connected to its lineage (kingdom [here: domain], phylum, class, order, family, genus, and species) in the “microbiomeanalyst_tax.txt” file. Taxa with an empty name were called “NoName” in the file and “UNMATCHED” taxa were set to “NA” (Section 3.1). Sample metadata were provided in the “microbiomeanalyst_meta.txt” file and linked to the lineage counts in the “*_otu.txt” file using the unique sample keys. Exported metadata included values such as antibiotics and probiotics usage, the sampling time point, and the category of the weight-for-age z-scores (Supplementary Table S5). Missing values in the metadata were replaced by “NA.”

### 3.4 Web-based interface

The web interface for MetagenomicsDB (https://www.bioinformatics.uni-muenster.de/tools/metagenomicsDB; accessed 2024/01/22) enabled our team members, regardless of bioinformatics expertise, to store, explore, and export the preliminary study results. The web interface was conceptually simple, as most of the data for the display are already aggregated in the views and the import and export are performed by the respective components of MetagenomicsDB (Sections 3.1 - 3.3). The only exception is the export of sequences in FASTQ format. The web interface was built based on the concept of tabs. Tabs contain collections of related data and are available for patients, samples, and taxa. The patients tab contains the patient metadata, selected static measurements (Supplementary Tables S3 and S5), and derived measurements related to the mother (Supplementary Tables S4 and S5). Samples and sequence counts per sample, selected variable measurements (Supplementary Table S3), and z-score (sub-)categories (Supplementary Table S4) are shown in the samples tab (Supplementary Table S5). Here, users can also export sequences in FASTQ format for samples of interest. For performance reasons, the download is limited to at most to 200,000 sequences.

By selecting samples or patients (in this case, internally the associated samples were used) from the respective tabs, users can export data to MicrobiomeAnalyst. In an overlay, the default settings of the export can be adjusted (Section 3.3). The ZIP archive with the exported data is then downloaded to the user’s device. A new tab with the aforementioned “Marker Data Profiling” workflow is automatically opened in the user’s browser, so that files can be directly uploaded after the archive has been extracted. The extracted files are uploaded to the “Text table format” tab on the website of the “Marker Data Profiling” workflow (Figure 3). The file “microbiomeanalyst_otu.txt” is uploaded as “OTU/ASV table” (the associated check boxes are left unchecked), “microbiomeanalyst_meta.txt” as “Metadata file”, and “microbiomeanalyst_tax.txt” as “Taxonomy table” (Figure 3). For classifications based on RDP, “Not Specific/Other” has to be chosen for “Taxonomy labels” (Figure 3).

**Figure 3:**
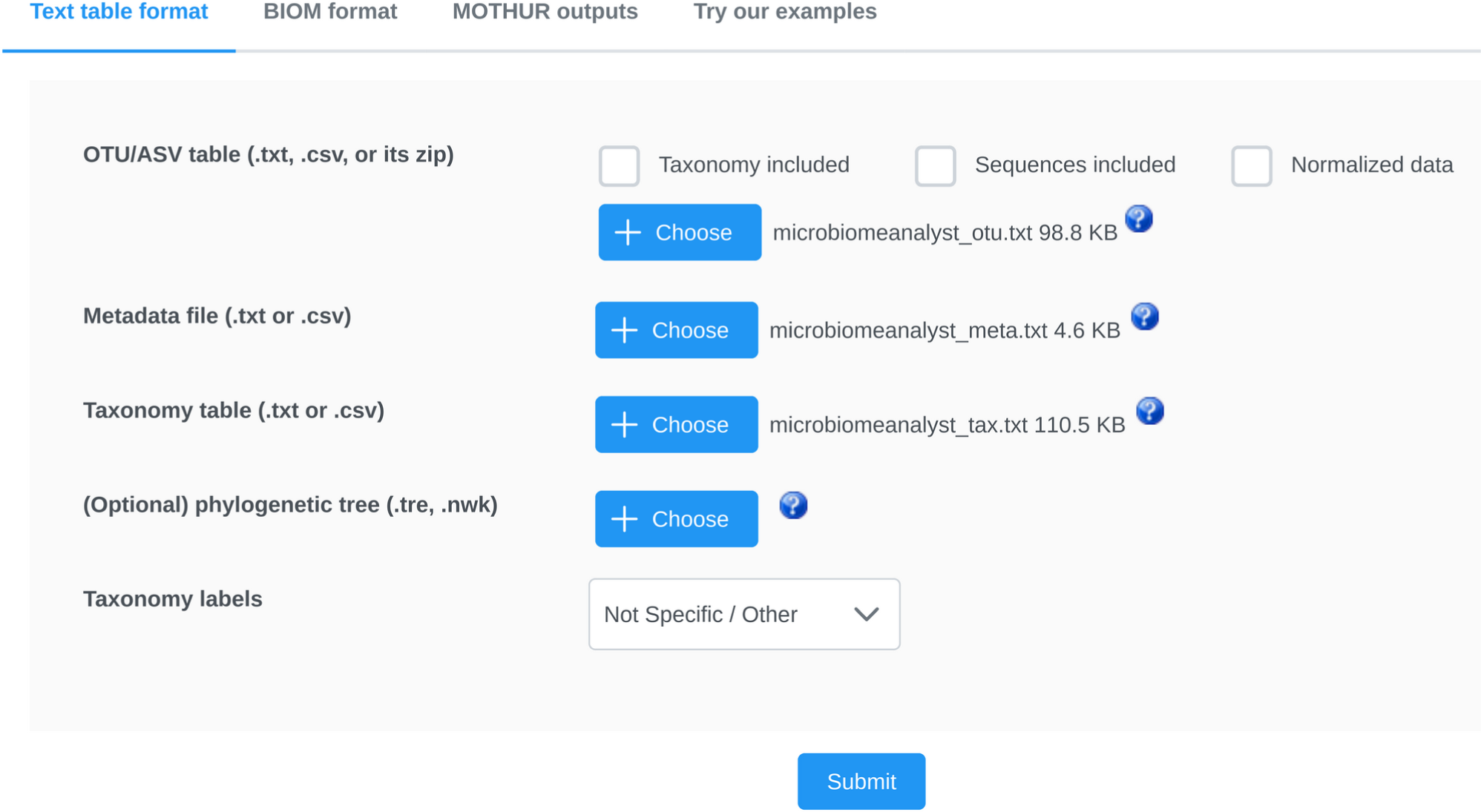
Upload interface for the “Marker Data Profiling“ workflow. This screenshot of the upload interface on MicrobiomeAnalyst (Lu et al. 2023) website (https://www.microbiomeanalyst.ca/MicrobiomeAnalyst/upload/OtuUploadView.xhtml; accessed 2024/02/01) illustrates how the files exported from MetagenomicsDB should be uploaded.

The taxa tab of MetagenomicsDB contains information about the taxa identified in each sample, the associated analysis program and database, and statistics on the supporting sequences (Section 3.2; Supplementary Table S5). Its display is limited to 10,000 entries in order to ensure good performance for users. Taxa in the taxa tab are grouped by name and rank and not by lineage in order to calculate the counts of supporting reads. In the RDP database that is used to classify the sequences (Supplemental Methods 4.2), taxa with identical names and the same rank can occasionally come from different lineages (Section 3.2). In these cases, read counts supporting the common taxon name can still be summed up. Users can click the question mark button in the “Taxon Name” field to assess, if taxa from different lineages were merged (Supplementary Table S5).

Using the name of the taxon, the tab provides several links to public resources which can be useful to obtaining further information, also with regard to potential health consequences of the observed taxon (Supplementary Table S5). We created a direct link to PubMed (Sayers et al. 2024) which queries the database for literature on the specific taxon with a focus on birth weight or SGA. The exact search term is given below. The position of the taxon name is indicated by the bold question mark:

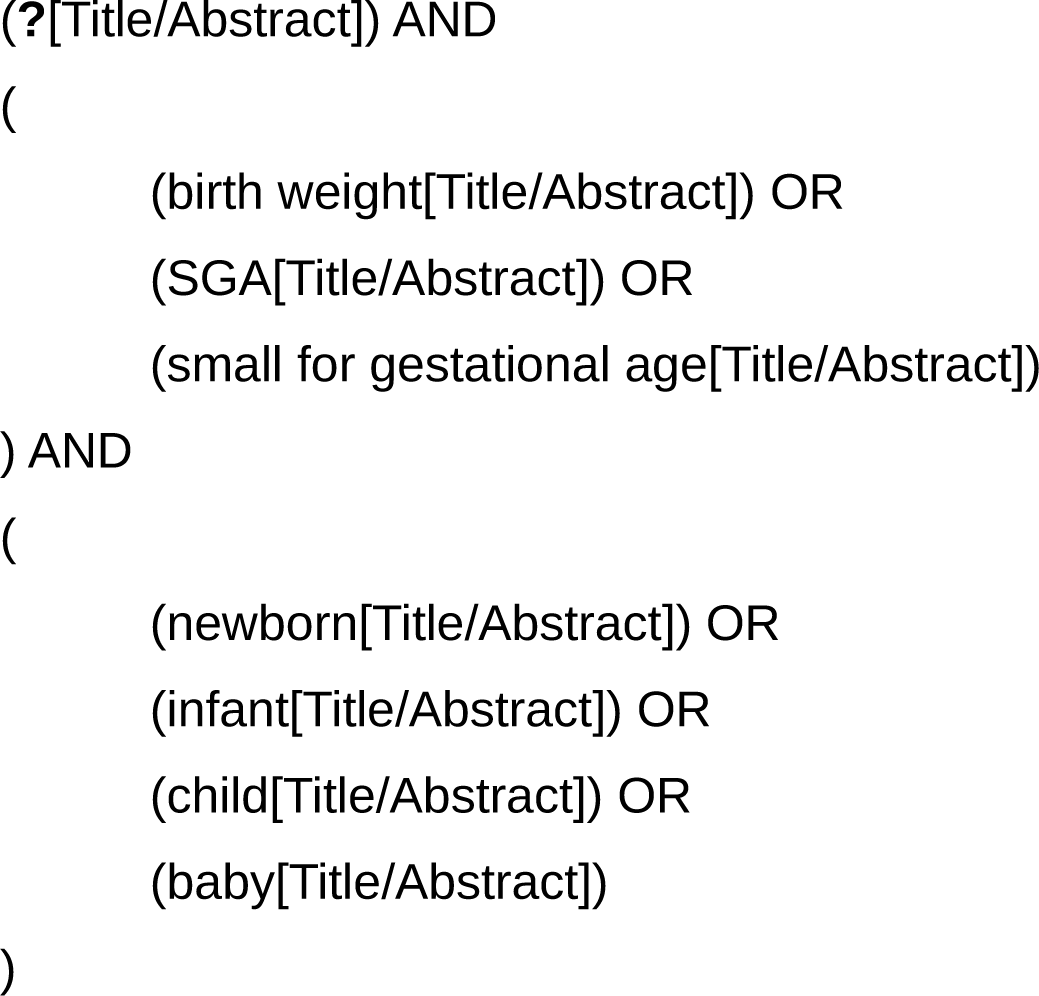

The interface also creates links to articles for the respective taxon on three other public resources: Wikipedia (https://en.wikipedia.org/; accessed 2024/02/02), the List of Prokaryotic names with Standing in Nomenclature (LPSN) (Parte et al. 2020), and the Bacterial and Viral Bioinformatics Resource Center (BV-BRC) (Olson et al. 2023), which are used to obtain information on the potential pathogenicity of the taxon. Links were created without *a priori* information, if the entry existed in the respective web resource. Thus, links might show no search results for certain taxa. However, we exclude special taxa like “UNMATCHED,” “FILTERED,” and “NA” (used for empty taxon name), since they are not expected to produce meaningful results (Section 3.1).

From the start page of MetagenomicsDB (Figure 4), tabs can be queried by selecting a tab of interest before entering search terms or making selections for searchable fields. It is not possible to conduct a search based on criteria from different tabs at the same time. Nevertheless, the results can be later refined in order to realize more complex queries (see below). Internally, searches in multiple fields are connected with an “and.” A letter in front of the search bars indicates the expected data type of the search term (Figure 4): “F” for floating point numbers, “I” for integers, and “T” for text. Advanced searches can be conducted by entering symbols together with the search term.

**Figure 4:**
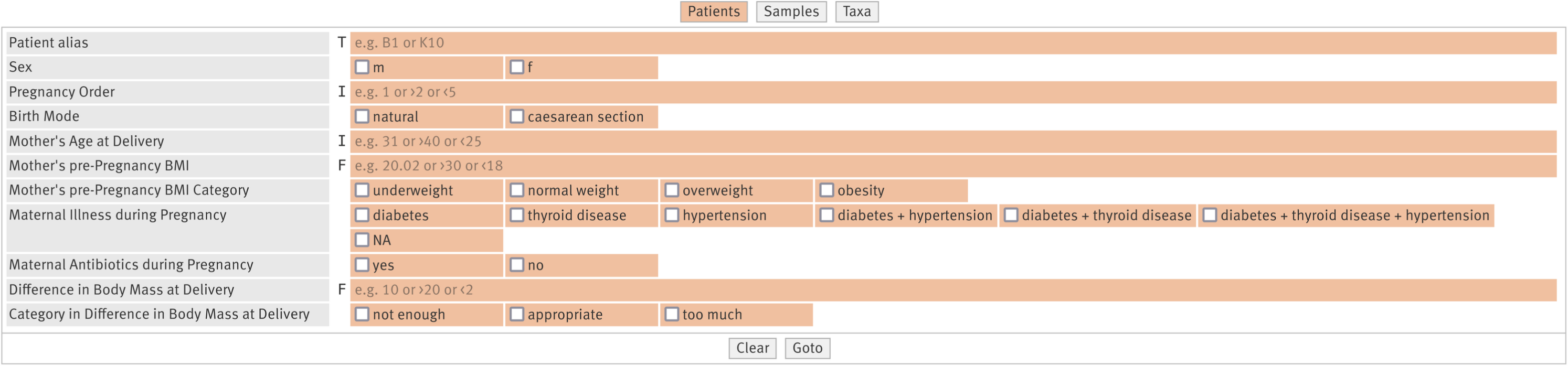
The search interface of MetagenomicsDB. The interface shows the search bars and checkboxes for searchable fields in the patients tab (https://www.bioinformatics.uni-muenster.de/tools/metagenomicsDB; accessed 2024/02/06). The letters in front of the search bars indicate the expected data types of the search terms: “T” for text, “I” for integers, and “F” for floating point numbers.

The mathematical signs “>=,” “<=,” “>,” and “<” can be used either individually or in combination (internally: “and”) to search integers and floating point numbers. By default, text-based searches have to have a full match and searches are case-sensitive. However, the use of “∼” triggers a pattern match. For example, entering “∼Escherichia” in the search bar for “Taxon Name,” searches for all names containing the term “Escherichia” (not case-sensitive). Multiple search terms can be joined by commas (internally: “or”). Likewise, if multiple checkboxes for a given field are selected, they are internally connected by an “or.” Users can also browse through a tab by submitting an empty query.

Once the user submits the search request, all database IDs matching the query are internally collected. Based on the IDs, all tabs are filled with data. Only the chosen tab is displayed, the remaining tabs are stored in the cache. When the user switches tabs, the currently displayed tab is exchanged with the selected tab from the cache. This approach does not require further database requests and thus reduces the loading time between tabs.

Search results can be further refined by selecting records of interest and clicking the “Apply” button. From hereon we will refer to this action as “applying a selection” in order to distinguish it from a different type of refinement (see below). When a selection is applied, the database IDs stored in the browser are replaced by the IDs of the selected records. This affects the remaining tabs as well. The operation can be undone using the “Undo” button. In contrast, the display of the records within the active tab can also be changed without affecting the internal selection of database IDs and thus, the other tabs. In the samples tab, the display of samples can be filtered by a minimum and/or maximum number of sequences (“# Sequences”). The display of records in the taxa tab can be filtered by the rank (“Ranks”), the average read length (minimum and maximum; “Avg. Read Length”), the average read quality (minimum and maximum; “Avg. Read Quality”), the supporting read count (minimum and maximum; “Count”), or any combination thereof. Subsequently, a new selection can be applied to the remainder of displayed records (see above). The combination of filtering the display in the active tab and of applying a selection enables more complex queries. To demonstrate this, we assessed how many patients had at least 5,000 reads with a minimum average quality of 10 supporting *Escherichia coli* in their “meconium” samples. We first searched for “Escherichia coli” in the “Taxon Name” field of the taxa tab on the main query page. Before a selection was applied to the remaining records, the display in the taxa tab was filtered by a minimum average read quality of 10 and a minimum read count of 5,000. In the samples tab, a selection was applied to the “meconium” samples. After switching to the patients tab, it became apparent that two patients (K19 and K27) matched this complex query.

## 4 Discussion

As of today, several programs are available which integrate data management and data analyses in a graphical interface. MG-RAST (Wilke et al. 2016) and MGnify (Richardson et al. 2023) both offer standardized analyses of sequences with associated metadata, data storage, and comparative analyses in a graphical web interface (Meyer et al. 2008; Wilke et al. 2016; Richardson et al. 2023). Both services offer to make results accessible to the public in their associated repositories (Meyer et al. 2008; Wilke et al. 2016; Richardson et al. 2023). The standardized workflows allow the comparison of results across studies in meta-analyses (Wilke et al. 2016; Richardson et al. 2023). In our study, our main goal is to store, explore, and analyze our own study data using an accessible interface. Thus, we chose a more flexible approach than permitted by the standardized workflows of both tools. First, we are interested in using the MetaG software for the classification of our 16S rRNA gene nanopore reads, since it has shown better performance on nanopore amplicons than other established programs (Manske, Grundmann, and Makałowski 2020). Secondly, our derived measurements contain dependencies on other measurement values and sometimes on external resources (as in the case of the weight-for-age z-scores), thus they are best calculated by a data management system based on stored data in order to maintain data integrity throughout the project (Section 3.1). Lastly, our custom data management system enables us to link our preliminary study results directly to more specialized web resources (Section 3.4).

Thus, we developed the central data management system MetagenomicsDB to help us store, explore, and analyze the data produced by our ongoing study. We argue that a central data management system can alleviate many of the challenges inherent to a microbiome study of a medical condition much better than conventional analyses based on data scattered over separate files. By bundling all preliminary results in a central location, MetagenomicsDB makes accessing the data straightforward, especially during a collaborative project. In our study, clinical, *in vitro*, and *in silico* analyses produced three distinct file types (Section 2). We needed to ensure the integrity of the data of each file type and maintain the correct connections between records in the different files. Using a more conventional approach with plain files, this would have been a tedious and error-prone manual task. Instead, we automated quality control in MetagenomicsDB using the general import unit (Figure 1), which parsed the inputs. By comparing the read IDs, for instance, it verified that the FASTQ files and the respective classification files were actually matching. It then inserted the quality-checked data into the storage component PostgreSQL (Figure 1). The automation of quality control helped to reduce human error, as opposed to the manual approach. Since transactions in PostgreSQL are ACID-compliant (https://www.postgresql.org/about/; accessed 2024/03/20), storing data in a PostgreSQL database instead of plain files had additional quality assurance advantages. The acronym ACID stands for atomicity, consistency, isolation, and durability (Haerder and Reuter 1983) and its principles comprise the following attributes with respect to changes of the stored data. Each transaction is carried out completely or not at all, thus avoiding partial updates (atomicity) (Gray 1981). In MetagenomicsDB, any synchronization between the database and the input data is conducted in a single transaction. Thus, even if just a single record is invalid, the synchronization can be completely undone, leaving the database in a consistent state. Changes to the data content must adhere to the constraints of the database management system (consistency) (Gray 1981). In the context of our system, this principle enables us to control the data types of the entered data, avoid duplicates, and maintain the connections between our different data types. According to the ACID principles, each transaction is carried out in isolation, despite concurrent activity (Meier and Kaufmann 2016, 136–38). The benefit for our collaborative project is that multiple investigators can work with the system at the same time without running the risk of overwriting each others changes. Once a transaction successfully modifies the data, the modified data are protected by the database management system from accidental loss (durability) (Gray 1981).

MetagenomicsDB is supplemented with a web-based graphical interface (Figure 1), which allows all investigators, regardless of bioinformatics expertise, to synchronize the study data with the database, explore, optionally filter the data using complex queries, and export them for statistical analyses in MicrobiomeAnalyst (Section 3.4). Thus, all team members receive live feedback on the progress of the study, which allows them to plan potential extensions, such as reanalyses or follow-up analyses, more efficiently. Furthermore, basic bioinformatic and statistical analyses of the preliminary study results are available to all team members. In order to further streamline the analysis workflow, we linked MetagenomicsDB to several public resources (Section 3.4). Wikipedia and LPSN provide a quick overview of the identified taxa. BV-BRC helps to assess the potential pathogenicity and thus the potential health impact of the microorganisms. Our customized PubMed search allows investigators to find information on already known links between the taxon of interest and SGA children more quickly. The web interface is not only expected to be useful for our internal purposes. By the time our study will be published, we aim to make the full study data available in MetagenomicsDB to reviewers and to the general public. Openly sharing the study results in an accessible way contributes to more transparent research and allows others to conduct follow-up analyses based on our findings. We demonstrated the viability of this approach by making a subset of our preliminary study results already publicly accessible: https://www.bioinformatics.uni-muenster.de/tools/metagenomicsDB

Despite the aforementioned advantages of a central data management system, if one was to develop such a system form scratch for every project, the investment in development time can be prohibitive. Thus, we designed MetagenomicsDB as a non-disposable data management system. Many components can be reused for future studies. The following design decisions contribute to an increased portability: first, the use of a relational database with a flexible schema (Section 3.2) which allows, among others things, to assign any number of measurements to a sample or to classify sequences with multiple programs or databases. Second, we export essential login information for the database to the standardized “.pg_service.conf” configuration file (Supplementary Methods 5.3.2), which helps to migrate our scripts to different computational environments. Third, we use a modular concept supplemented with tests for the import and export unit of MetagenomicsDB. Thus, several general functions (e.g. extracting files or parsing FASTQs and classification files) can be easily reused for future projects. At the time of writing, these functions are already flexible enough to support input file types (Supplementary Table S6) which are not currently used in this project. Certain high level-functions and features of the database schema necessarily are project-specific. Examples include the calculation of the weight-for-age z-scores and the assignment of patients to the weight-for-age z-score (sub-)categories. The web interface is also specific to our web environment. Nevertheless, due to the modular concept, project-specific functions can be replaced or adapted to new projects (Section 3) in a fraction of the time needed for developing a new data management system from scratch.

In conclusion, we have developed a data management system with an accessible web interface for the study of the gut microbiome in SGA children. We have outlined, how the system enhances quality assurance of the produced results, how it streamlines the *in silico* analyses in a collaborative project, and how it will be able to contribute to more transparent sharing of results at publication time. We acknowledge that despite the efforts towards portability, our system is not yet ready to be widely applicable out-of-the-box for any microbiome study. A major improvement in this direction is the development of a portable graphical interface. However, we expect that due to the modular concept, the tests, and the extensive description of the underlying rationale, large parts of our implementation can be adapted for future projects. Thus, we are reporting our endeavors to motivate the application of data management systems at the scale of single studies in microbiome research.

## 5 Methods

### 5.1 Study of the gut microbiome in small for gestational age children

We conducted a blinded prospective study on the potential influence of the gut microbiome on birth weight and weight gain in SGA children during the first year of life. The study cohort consisted of otherwise healthy (Supplementary Methods 1) single birth children of Polish descent and was recruited from the Heliodor Święcicki Clinical Gynecology and Obstetrics Hospital of Poznań University of Medical Sciences (Poznań, Poland). No siblings were present in the cohort and all children were born on time (Supplementary Methods 1). Newborns were divided into cohorts of appropriate for gestational age and SGA children (Supplementary Methods 1). 65 children were examined in regular intervals (Supplementary Methods 1) from birth (“meconium”) to the age of 1 year (“1y”) in order to assess the weight gain and to sample the gut microbiome. Samples were skipped if the children were hospitalized or had diarrhea when the sample was scheduled. During each appointment, children were weighted and stool samples were collected (Supplementary Methods 2 and 3.1). Patient and maternal metadata, measurement results, confounding factors to the gut microbiome (antibiotics, probiotics, and feeding habits), and selected data on the course of the pregnancy, including maternal health, were stored in an Excel spreadsheet together with derived measurements (Supplementary Methods 2; Supplementary Table S1). Derived measurements represented values which were calculated based on other values in the spreadsheet (e.g. “mother’s age at delivery”, Supplementary Table S1).

### 5.2 Sequencing of stool samples and *in silico* analyses

In the lab, DNA was extracted from the fecal samples using the DNeasy PowerSoil Pro Kit (QIAGEN GmbH, Hilden, Germany) (Supplementary Methods 3.2). After verifying that the extraction was successful (Supplementary Methods 3.2), we created full-length 16S rRNA gene amplicons using the 16S Barcoding Kit 1-24 (Cat. No.: SQK-16S024, Oxford Nanopore Technologies plc, Oxford, England). 23 of the 24 available barcodes were used for the genuine samples. The remaining barcode (by our definition: 99, Supplementary Methods 5.3.1) was assigned to a sample containing distilled water, which served as an internal negative control for library preparation and sequencing (Supplementary Methods 3.3). The finished library was sequenced for 24 hours using two MinION Mk1B nanopore sequencers (Cat. No.: MIN-101B, Oxford Nanopore Technologies plc, Oxford, England) with Flongle Flow Cells (Cat. No.: FLO-FLG001, Oxford Nanopore Technologies plc, Oxford, England). Real-time basecalling was performed using Dorado v0.3.3 in MinKNOW version 23.07.5 (https://community.nanoporetech.com/downloads; accessed 2024/06/08) (Supplementary Methods 3.3). After manual quality control (Supplementary Methods 3.3), FASTQ “pass” files were processed with MetaG. We removed residual human contamination (Supplementary Methods 4.1) before classifying the remainder using RDP version 11.5 (Supplementary Methods 4.2). Both analyses used new MetaG standard parameters, which were generally applicable and not specific to the study of the gut microbiome (Supplementary Methods 4.1.1, 4.1.2, 4.2.1, and 4.2.2). FASTQ “pass” files and MetaG classifications were stored in a common directory with subdirectories named after the run and the barcode of the 16S rRNA gene reads (Supplementary Methods 5.3.1). The run and the barcode from sequencing the stool samples, the classification program (MetaG), and the classification database (RDP) were added to the Excel spreadsheet (Supplementary Table S1). We used the run and the barcode as a unique key to assign the samples from the spreadsheet to the respective sequences and classifications.

### 5.3 Development of a central data management system

We developed a central data management system called MetagenomicsDB to allow all users, regardless of bioinformatics expertise, to store, explore, and export their preliminary study results (Sections 5.1 and 5.2) in a format suitable for downstream statistical analyses. During development, we aimed to create a modular system which could be adapted for future microbiome studies. MetagenomicsDB consists of three main units (Figure 1) for import and quality control, storage, and export. Our system is supplemented with an accessible web interface, which is hosted on an Apache (https://apache.org/; accessed 2024/03/22) 2.4.48 server running under FreeBSD 13 (https://www.freebsd.org/; accessed 2024/03/17). It uses in-house JavaScript and Cascading Style Sheets. The interactions between the web interface and the users are protected by Hypertext Transfer Protocol Secure (HTTPS). The import and export unit, as well as the helper scripts for the web interface, are implemented in Perl 5. Preliminary study results and weight-for-age growth standards from the WHO (https://cdn.who.int/media/docs/default-source/child-growth/child-growth-standards/indicators/weight-for-age/expanded-tables/wfa-boys-zscore-expanded-tables.xlsx and https://cdn.who.int/media/docs/default-source/ child-growth/child-growth-standards/indicators/weight-for-age/expanded-tables/wfa-girls-zscore-expanded-tables.xlsx; accessed 2024/01/30) are stored in a relational PostgreSQL 14beta2 database located on a database server with two Intel E5-2660 v2 CPUs, 384 GB RAM, and three 8 TB hard drives running under FreeBSD 13. Our data management system is described in depth in Section 3 and Supplementary Methods 5.

## 6 Ethics

The experimental protocol was established in accordance with the ethical guidelines of the Declaration of Helsinki and approved by the Bioethics Committee of the Poznań University of Medical Sciences in Poznań, Poland (Resolution No. 248/20). Since the study includes children, written informed consent was obtained from their parents or legal guardians before entering the study and after the aims and methodologies of the study were explained.

## 7 Availability

The source code for MetagenomicsDB is available from our GitHub repository (https://github.com/IOB-Muenster/MetagenomicsDB; accessed 2024/06/07). The MetaG source code can be found at https://github.com/IOB-Muenster/MetaG (accessed 2024/06/07).

## Acknowledgments

We wish to thank Michelle Leyers for her help with the classification of sequences using the MetaG software and Marc-Nicolas Bendisch who contributed to the benchmarking of the MetaG, results which was the basis for the updated parameter training in MetaG. Parts of the calculations for this publication were performed on the High Performance Computing (HPC) cluster Paralleles Linux-System für Münsteraner Anwender (PALMA) II of the University of Münster, subsidized by the German Research Foundation (INST 211/667-1).

## Supplementary Information for

### Supplementary Methods

#### Availability

The MetagenomicsDB source code is available from our GitHub repository (https://github.com/IOB-Muenster/MetagenomicsDB; accessed 2024/06/07). The MetaG source code can be found at https://github.com/IOB-Muenster/MetaG (accessed 2024/06/07). In the following, the major steps of the computational analyses are illustrated by command line examples. For the sake of readability, we left out parameters purely related to parallelization and to environment variables. Both have to be adjusted specifically to the reader’s computing environment. More details can be found in the documentation of the scripts (for example the help function or help text in the scripts).

### 1 Study design

We conducted a blinded prospective study with single birth children of Polish descent. The aim was to investigate the possible influence of the gut microbiome on birth weight and subsequent weight gain in small for gestational age children (SGA) within the first year of life. No siblings were present in the study. Parents and their children were recruited from the Heliodor Święcicki Clinical Gynecology and Obstetrics Hospital of Poznań University of Medical Sciences (Poznań, Poland). Children were divided in cohorts (z-score category) in accordance with the eleventh revision of the World Health Organization’s (WHO’s) International Classification of Diseases and Related Health Problems (ICD-11) (World Health Organization 2022, 1216): newborns were classified as SGA if their weight-for-age z-scores at birth were less than negative two. The calculation of the z-scores is explained in more detail in Supplementary Methods 5.1.2. In our study, non-SGA children were assigned to the appropriate for gestational age (AGA) category. SGA and AGA children were hospitalized in different wards of the hospital: the Department of Neonatology and the Department of Obstetrics, respectively.

Newborns were only eligible for the study if they were born on time (37-42 weeks) (World Health Organization 2022, 1220,1222). The fetal age was determined by a neonatologist in accordance with the Naegele rule (Law and Martin 2020, 380), corrected by an ultrasound assessment in the first trimester of pregnancy. Patients had to be healthy, apart from the condition under study (SGA vs. AGA). To ensure this, we evaluated clinically diagnosed risk factors for abnormal pregnancy, prenatal development disorders, and maternal health during pregnancy. Namely we considered: viral, bacterial, and parasitic infections; vaginal infections; intrauterine infections, such as intrauterine pneumonia; placental inflammation; disturbances of the fetal water volume; irregularities of the placenta; premature rupture of the fetal membranes; previous miscarriages; hypoxia; and perinatal outcome, APGAR score (Apgar 1953), and length of hospitalization of the newborns. Children with diagnosed hypoxia, congenital heart disease, neural tube malformations, genetic defects, and diseases requiring surgery were excluded from the study. After applying the exclusion criteria, 65 patients remained, some of whom withdrew throughout the course of the study.

The remaining children were examined at up to eight time points in order to assess the weight gain (Supplementary Methods 2) and gut microbiome (Supplementary Methods 3) during the development: at birth (“meconium”), after three days (“3d”), two weeks (“2w”), six weeks (“6w”), three month (“3m”), six month (“6m”), nine month (“9m”), and one year (“1y”). Definitions of time points were fuzzy to account for minor variation in the sample collection date (Supplementary Methods 5.1.2). Sampling time points were skipped if children were hospitalized or suffered from diarrhea.

### 2 Measurements of patients and their mothers

For the first two samples (Supplementary Methods 1), the body mass of the children was recorded by the medical staff at the Poznań University of Medical Sciences (Poznań, Poland) using a High-Resolution Digital Neonatal Four-Sided Tray Scale (Cat. No.: 2210KL4CT, Health o meter, McCook, USA) with an accuracy of 1 g in the eligible range of birth weights (Supplementary Methods 1). For the remaining time points (Supplementary Methods 1), weights were recorded by pediatricians using medical scales with an accuracy of 100 g. Pediatricians from the Department of Neonatology (Poznań University of Medical Sciences) collected data on the course of the pregnancy, the mother’s health status, and the newborns based on the maternity record, interviews with the mothers, and neonatal discharge cards. This included information on confounding factors to the gut microbiome: antibiotics and probiotics use and feeding habits. We stored the relevant raw data from pediatricians in a spreadsheet and added further measurements which we calculated based on the raw data (Supplementary Table S1). This file also contained a key (“number of run and barcode”), which linked the spreadsheet to the sequencing and classification results from analyzed stool samples (Supplementary Methods 3.3, 4.2, and 5.3.1). Data which we used to ensure that patients were healthy apart from the condition under study (Supplementary Methods 1) were not recorded in the spreadsheet.

**Table S1:** Columns in the measurement Excel file. Definition of columns in the measurement Excel file. Grey shading is used for columns which were calculated based on other columns.

| Column name | Definition |
| --- | --- |
| patient ID | Pseudonym of the patient in the study. |
| hospital code | Patient's accession number from the hospital. |
| birth date | Patient's birthdate. |
| sex | Sex of the patient:<br>• "1" for "m"<br>• "2" for "f" |
| time point | Text string describing the "sample collection date" relative to the "birth date" (Supplementary Methods 5.1.2). |
| sample collection date | Date when the sample was taken. |
| body mass | Body mass of the patient (g). |
| body mass z-score | Weight-for-age z-score (Supplementary Methods 5.1.2). |
| catch-up | Obsolete rating of the "body mass z-score":<br>• "1" for "no catch-up"<br>• "2" for "catch-up"<br>• "3" for "AGA child"<br>Current ratings are presented in Supplementary Methods 5.1.2. |
| birth mode | How the patient was born:<br>• "1" for "natural"<br>• "2" for "caesarean section" |
| feeding mode | How the patient was fed:<br>• "1" for "breastfed"<br>• "2" for "formula"<br>• "3" for "mixed"<br>• "4" for "diet extension" |
| probiotics | Did the patient receive probiotics?<br>• "1" for "yes"<br>• "2" for "no". |
| antibiotics | Did the patient receive antibiotics?<br>• "1" for "yes"<br>• "2" for "no" |
| mother's birth date | The birthdate of the patient's mother. |

**Table S1: Columns in the measurement Excel file (continued).**
|  |  |
| --- | --- |
| mother's age at delivery | The age of the mother at the patient's birth (years) (Supplementary Methods 5.1.2). |
| maternal body mass before pregnancy | The mother's weight before getting pregnant with the patient (kg). |
| maternal body mass at delivery | The mother's weight when giving birth to the patient (kg). |
| mother's height | The height of the mother (m). |
| mother's pre-pregnancy BMI | The body mass index (BMI) (Keys et al. 1972) of the mother before getting pregnant with the patient (Supplementary Methods 5.1.2). |
| pre-pregnancy BMI category | Rating of "mother's pre-pregnancy BMI" (Supplementary Methods 5.1.2). |
| difference in body mass at delivery | Difference between "maternal body mass at delivery" and "maternal body mass before pregnancy" (Supplementary Methods 5.1.2). |
| category of difference in body mass at delivery | Rating of "difference in body mass at delivery" (Supplementary Methods 5.1.2). |
| pregnancy order | The patient is the mother's n-th child where n is the pregnancy order. |
| maternal illness during pregnancy | Mother's illness while being pregnant with the patient:<br>• "1" for "diabetes"<br>• "2" for "thyroid disease"<br>• "3" for "hypertension"<br>• "4" for "diabetes + thyroid disease"<br>• "5" for "diabetes + hypertension"<br>• "6" for "thyroid disease + hypertension"<br>• "7" for "diabetes + thyroid disease + hypertension" |
| maternal antibiotics during pregnancy | Did the mother receive antibiotics while being pregnant with the patient?<br>• "1" for "yes"<br>• "2" for "no" |
| program | The program that was used to classify the reads (Supplementary Methods 4.2). |
| database | The classification database that was used with the "program" to classify the reads (Supplementary Methods 4.2). |
| number of run and barcode | Run and barcode of the sequencing results for the collected stool sample. Used to connect this file with the FASTQs and MetaG results (Supplementary Methods 3.3, 4.2, and 5.3.1). |

### 3 Processing of the stool samples

#### 3.1 Sample collection

Meconium and fecal samples on the third day after delivery were collected by medical staff using professional swabs with a 1 mL eNAT Transport and Preservation Medium stabilizing the ribonucleic acid (RNA) and the deoxyribonucleic acid (DNA) of bacteria in the sample (Cat. No.: 608CS01R, Copan Italia S.p.A., Brescia, Italy). Samples were subsequently frozen at -20°C. After the children left the hospital, samples were collected by the parents or guardians in the time intervals specified in Supplementary Methods 1 using the aforementioned collection kit. Feces were obtained from the diapers according to a standardized procedure. All samples were transported to the Institute of Human Biology and Evolution at the Faculty of Biology (Adam Mickiewicz University, Poznań, Poland), where they were stored at -20°C before DNA extraction.

#### 3.2 DNA extraction

After suspending stool and meconium samples in stabilizing buffer (Section 3.1), 250 μL from each sample was used to extract the total genomic DNA using the DNeasy PowerSoil Pro Kit (QIAGEN GmbH, Hilden, Germany) according to the manufacturer’s instructions. Deviating from the instructions, we incubated samples for 10 minutes at 65°C before homogenization, so as to achieve better results during lysis (data not provided). Homogenization was performed using PowerBead Pro tubes (Cat. No.: 19301, QIAGEN GmbH, Hilden, Germany) and TissueLyser II (Cat. No.: 85300, QIAGEN GmbH, Hilden, Germany) set to a maximum oscillation frequency for a duration of five minutes. Each sample was then eluted in 50 μL of solution C6 from the DNA extraction kit. The solution was preheated to 60°C. After DNA isolation, 1 μL from each sample was used to quantify the DNA using a Qubit 4 Fluorometer (Thermo Fisher Scientific Inc., Waltham, Massachusetts, USA) to ensure that isolates had sufficient quality for the subsequent library preparation and sequencing. After this step, isolates were stored at -20°C.

#### 3.3 Full-length 16S ribosomal ribonucleic acid gene sequencing

Full-length 16S ribosomal ribonucleic acid (rRNA) gene sequence libraries were prepared with the 16S Barcoding Kit 1-24 (Cat. No.: SQK-16S024, Oxford Nanopore Technologies plc, Oxford, England) according to the manufacturer’s instructions using 10 ng of DNA from each sample. From the 24 available unique barcodes per run, up to 23 were used for the genuine samples. A water control sample containing Invitrogen UltraPure DNase/RNase-Free Distilled Water (Thermo Fisher Scientific Inc., Waltham, Massachusetts, USA) was included in each run using the remaining barcode (barcode 99 in Supplementary Methods 5.3.1) in order to check for contamination during library preparation and sequencing. The 16S rRNA gene sequences were amplified with the ProFlex 3 x 32-well PCR-System (Thermo Fisher Scientific Inc., Waltham, Massachusetts, USA) using the following program: 1 min denaturation at 95°C, 25 cycles (each: 95°C for 20 s, 55°C for 30 s, 68°C for 120 s) and a final extension step at 65°C for 5 minutes. Amplicons were then purified on magnetic beads (Agencourt AMPure XP, Beckman Coulter Inc., Brea, USA) and quantified using the aforementioned Qubit 4 Fluorometer. Equal amounts of amplicons from each sample were pooled and the library was processed according to the manufacturer’s recommendations. The library was subsequently sequenced for 24 hours using two MinION Mk1B nanopore sequencers (Cat. No.: MIN- 101B, Oxford Nanopore Technologies plc, Oxford, England) with Flongle Flow Cells (Cat. No.: FLO-FLG001, Oxford Nanopore Technologies plc, Oxford, England). Real-time basecalling was performed using Dorado v0.3.3 in MinKNOW version 23.07.5 (https://community.nanoporetech.com/downloads; accessed 2024/06/08) on a Windows 11 Pro 23H2 computer with an Intel Xeon W-2135 CPU, 32 GB RAM, and 2.5 TB of disk space. The following settings were applied: the basecalling mode was set to fast, modified basecalling was deactivated, and the voltage was set to 180 mV. Minimum barcode and read quality scores of 60 and 7, respectively, were applied. Reads were output in the FAST5 and FASTQ formats. We repeated library preparation and sequencing of the barcode or of the whole run if no FASTQ “pass” reads were output, the number of FASTQ “fail” reads exceeded that of FASTQ “pass” reads, or the barcode of a sample was not read. Only FASTQ “pass” files were used for downstream analysis (Supplementary Methods 4) and the run and barcode of non-control sequences were added to the Excel spreadsheet containing the patient and maternal metadata and the measurements (“number of run and barcode,” Supplementary Table S1).

### 4 *In silico* analysis of FASTQ files

After basecalling, residual human reads were removed from FASTQ “pass” files. The remainder was classified using MetaG (Manske, Grundmann, and Makałowski 2020) and the RDP database (Cole et al. 2014) (Supplementary Figure S1; Supplementary Methods 4.1 and 4.2). We made several adjustments to the MetaG core algorithm since its first publication in 2020 (Manske, Grundmann, and Makałowski 2020). The following sections will describe updates that were relevant for this study, including the aforementioned workflow to remove human reads (Supplementary Methods 4.1) and the generation of standard parameters which were used for the current analyses (Supplementary Methods 4.1.1, 4.1.2, 4.2.1, and 4.2.2).

**Figure S1:**
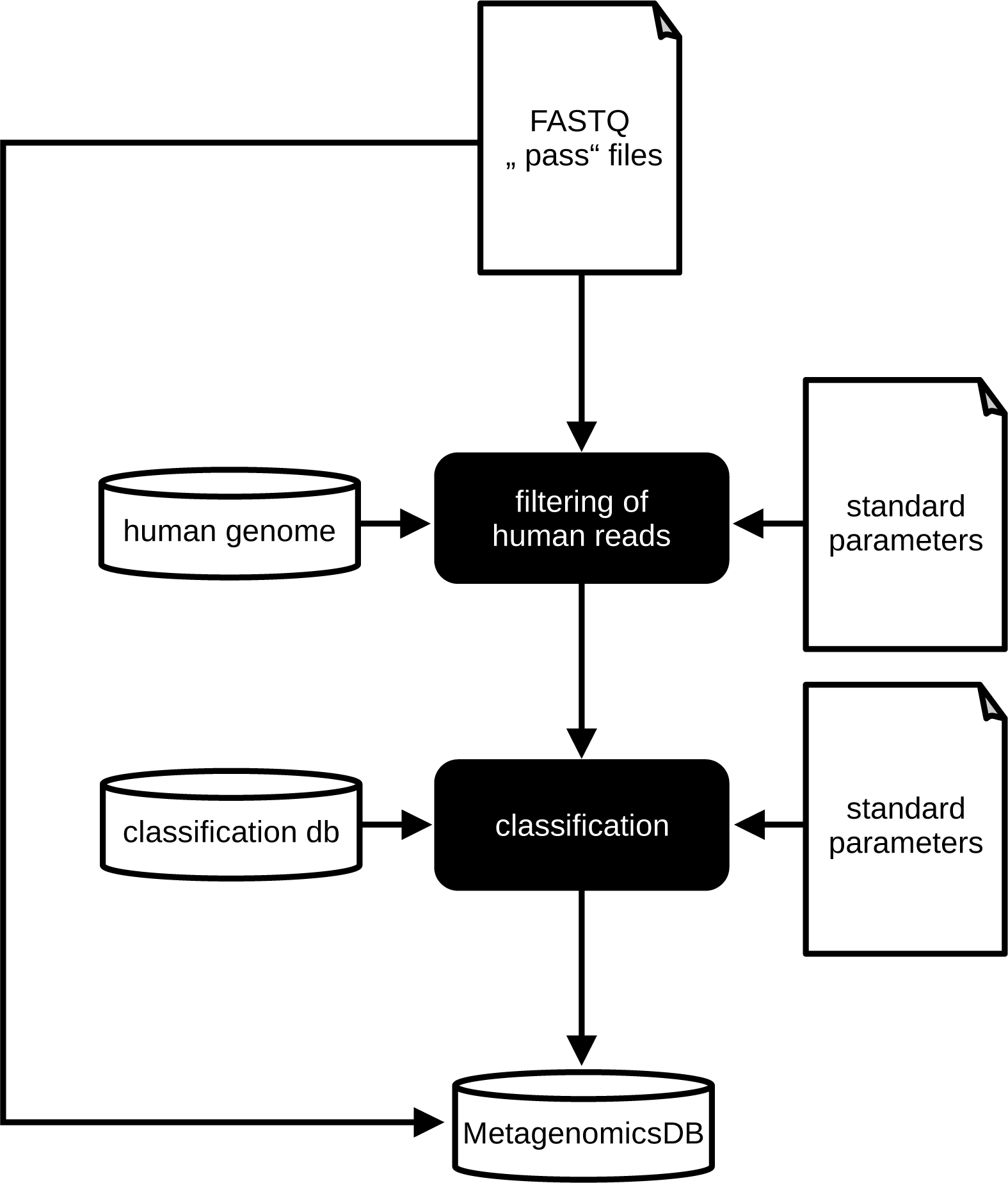
MetaG analysis workflow. MetaG (Manske, Grundmann, and Makałowski 2020) removed human reads from the FASTQ “pass” files (Supplementary Methods 3.3) using the T2T human reference genome (“human genome”) and the appropriate standard parameters (Supplementary Methods 4.1). The remaining reads were subject to taxonomic classification using the classification database (“classification db”) and its standard parameters (Supplementary Methods 4.2). Sequences and classifications were stored in MetagenomicsDB (Supplementary Methods 5). This figure was generated with LibreOffice Draw 7.0.4.2 (https://www.libreoffice.org/discover/draw/; accessed 2024/03/22).

#### 4.1 Removal of human reads

A major update to the MetaG algorithm since its first publication (Manske, Grundmann, and Makałowski 2020) was the addition of a filtering workflow which allowed users to remove contamination (commit 275c4a9; https://github.com/IOB-Muenster/MetaG; accessed 2024/01/22). This reduced false classifications by off-target reads and helped to protect the privacy of patients when human host contamination was a concern. The basic algorithm behind the filtering workflow was to perform a LAST (Kiełbasa et al. 2011) alignment between the input reads and the database containing the genome(s) of the organism(s) to be removed. Only reads that did not align to this filter database were extracted from the original input sequence file and written to an output FASTA file (Supplementary Figure S1). The filtered reads could then be used for follow-up taxonomic classifications (Supplementary Figure S1). Here, we built a database based on the T2T- CHM13v2.0 human reference genome (RefSeq (Sayers et al. 2024) accession: GCF_009914755.1). All nucleotides were transformed to an upper case to remove soft- masking of repeats before LASTdb (Frith, Hamada, and Horton 2010) version 1256 was used to build a database for MetaG.

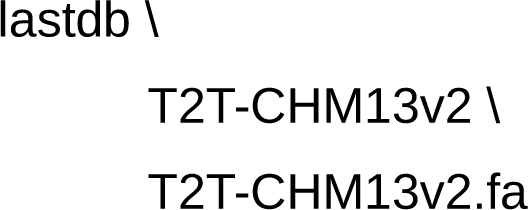

We used the local version of MetaG (commit 275c4a9; https://github.com/IOB-Muenster/MetaG; accessed 2024/01/22) on the Paralleles Linux-System für Münsteraner Anwender (PALMA) II High Performance Computing cluster of the University of Münster to remove residual human contamination from our samples (--filter) using a set of standard parameters (--config) and our newly built database (-ldb). Parameter training with the help of LAST-TRAIN (Hamada et al. 2017) and LAST-SPLIT (Frith and Kawaguchi 2015) is described in the following sections (Supplementary Methods 4.1.1 and 4.1.2; Supplementary Figure S2). All FASTQ files were located in a single input directory (-q) and were processed using the recursive mode (--recursive) of MetaG.

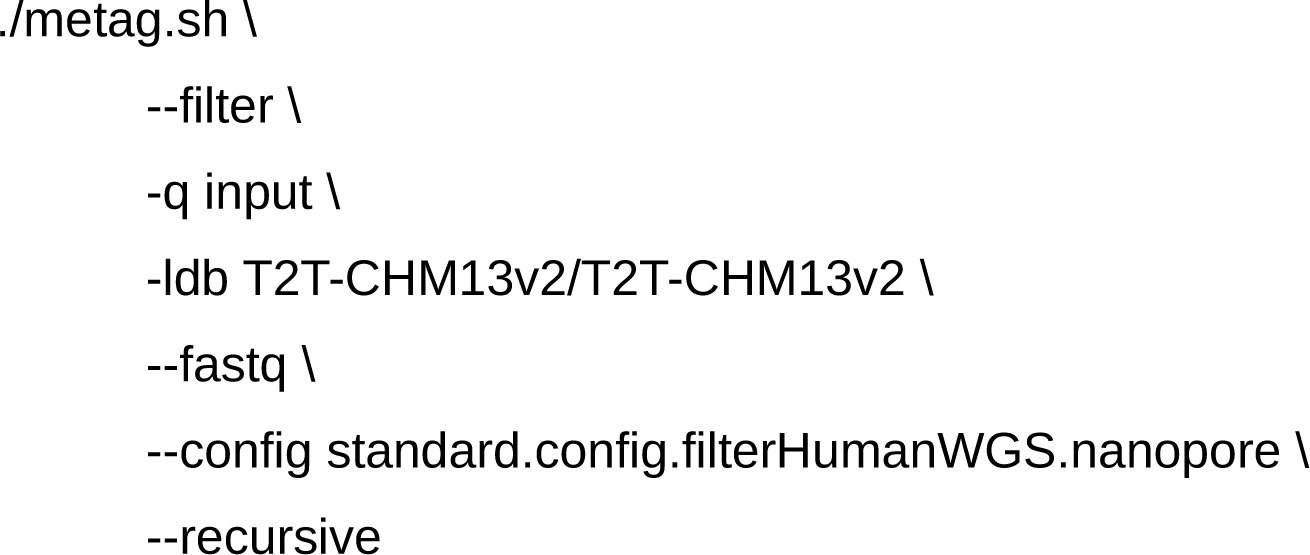

**Figure S2:**
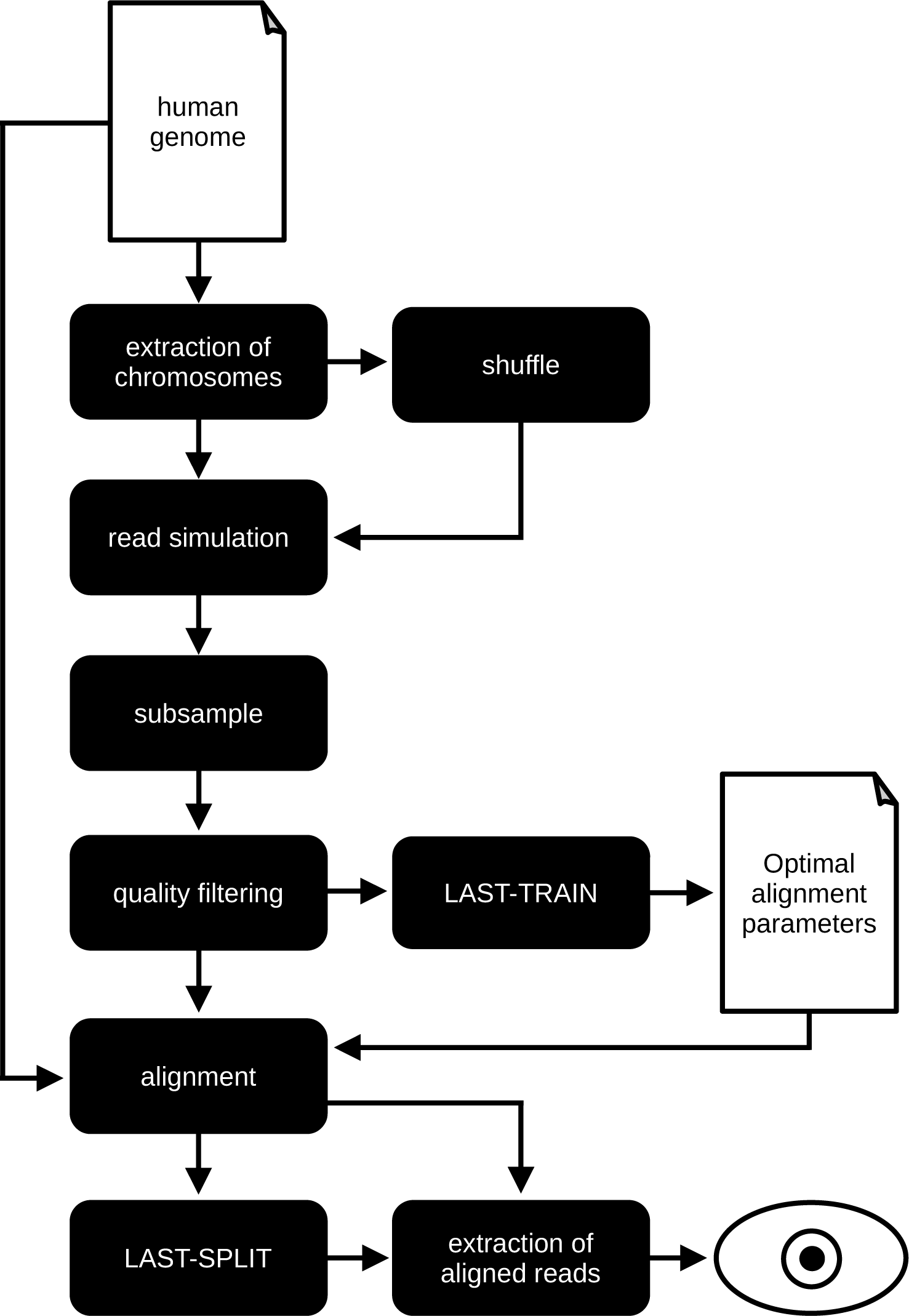
Workflow for the training of filter parameters. Chromosome sequences were extracted from the T2T human reference genome (“human genome”; Supplementary Methods 4.1) and a shuffled copy was generated from each sequence. All copies were used to simulate sequencing. Subsequently, reads were subsampled and filtered by quality (Supplementary Methods 4.1.1). Optimal alignment parameters were sought with LAST-TRAIN (Hamada et al. 2017) and used to generate an optimal alignment of the retained reads and the reference genome. The raw alignment was filtered by multiple runs of LAST-SPLIT (Frith and Kawaguchi 2015) (Supplementary Methods 4.1.2). Aligned reads were extracted from each alignment and manually compared (eye symbol) to assess if LAST-SPLIT filtering was necessary and if so, which settings were appropriate (Supplementary Methods 4.1.2). This figure was generated with LibreOffice Draw 7.0.4.2 (https://www.libreoffice.org/discover/draw/; accessed 2024/03/22).

##### 4.1.1 *In silico* simulation of human reads

In order to identify alignment parameters for the human T2T database which would ideally remove all human reads, but at the same time keep all non-human reads in the input, we simulated nanopore sequencing from the same T2T reference genome that had already been used for database building (Supplementary Methods 4.1). For this purpose, we split the FASTA file by chromosome (Supplementary Figure S2). An additional shuffled copy of each chromosome with preserved mononucleotide composition was generated (Supplementary Figure S2) using easel 0.48 (https://github.com/EddyRivasLab/easel; accessed 2022/01/25) from HMMER (Eddy 2011) 3.3.2. The shuffled copies were used as negative controls in order to ensure that filtering would not be overly strict. From each copy of each chromosome, we simulated nanopore sequencing (Supplementary Figure S2) with a target coverage of 2x (--quantity) using Badread (Wick 2019) v0.2.0 (as exemplified below for the raw copy of chromosome one). Using SeqKit (Shen et al. 2016) v2.0.0, 5% of reads from each genuine chromosome and 0.5% from each shuffled chromosome were retained (-p) in order to obtain more manageable training sets (Supplementary Figure S2). This is exemplified below for the raw copy of chromosome one.

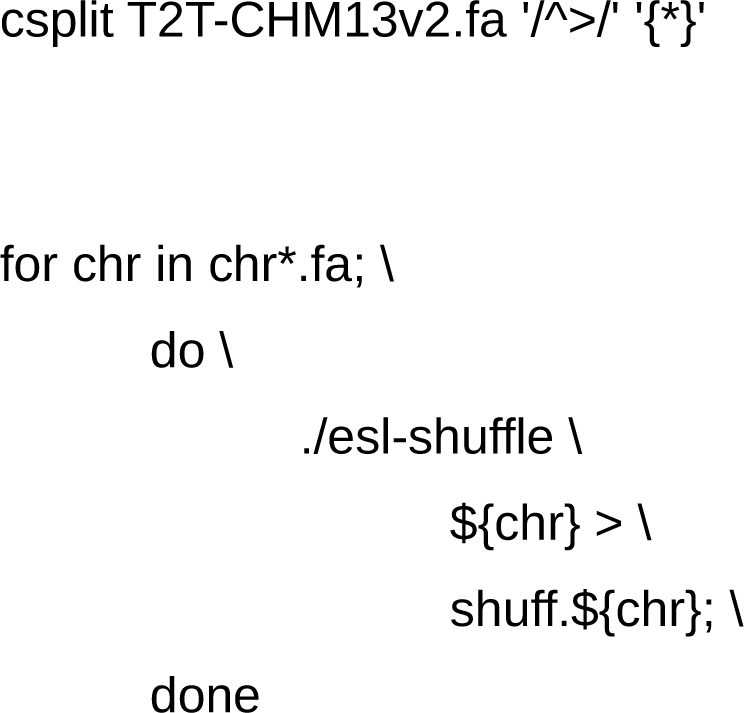

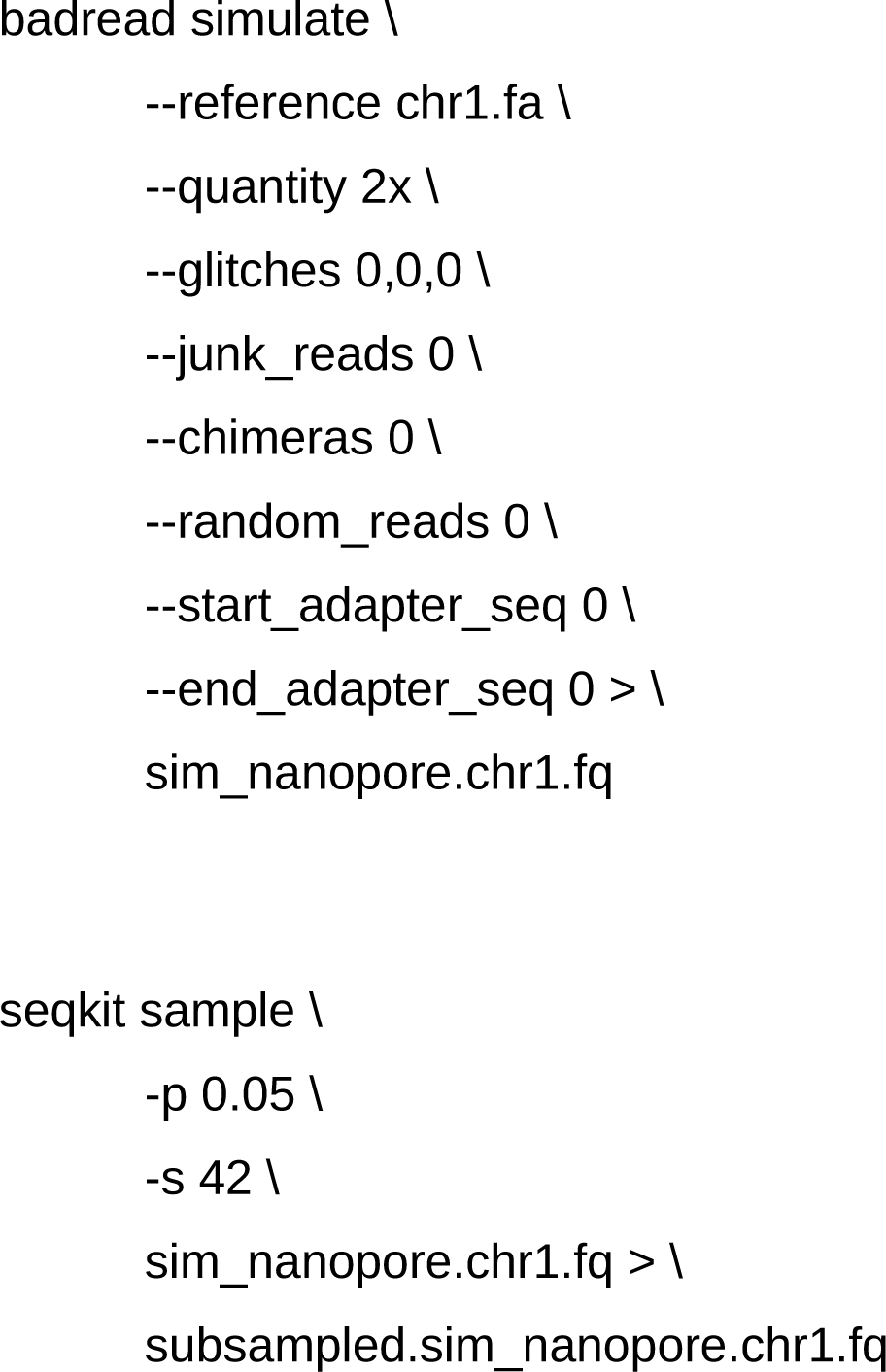

All simulated reads were pooled and subsequently subject to a quality control pipeline (Supplementary Figure S2). Our Badread simulations produced demultiplexed and already basecalled reads without any adapters or primers. According to the recently published literature on amplicon or metagenomic analysis using MinION, quality filtering using the basecaller’s defaults (Chng et al. 2020; Leggett et al. 2020) (we assumed usage of quality-filtered “pass” reads, if not explicitly stated otherwise) or by applying a relatively modest cutoff (average read quality score > 7-9) (Urban et al. 2021; Nygaard et al. 2020) were customary pre-processing steps before downstream analyses. The error profile of Badread in our simulation was based on a nanopore sequencing run from 2020. So assuming flowcell version FLO-MIN107 and a DNA sequencing kit such as SQK-LSK108 or SQK-LSK109 were used, the minimum quality of a “pass” read according to the default settings of the Guppy basecaller (https://community.nanoporetech.com/downloads; accessed 2022/02/07) version 6.0.1+652ffd1 was seven (configuration file: dna_r9.5_450bps.cfg). For this reason, we excluded any read with an average quality of less than seven (-q) from the simulated reads using NanoFilt (De Coster et al. 2018) version 2.8.0.

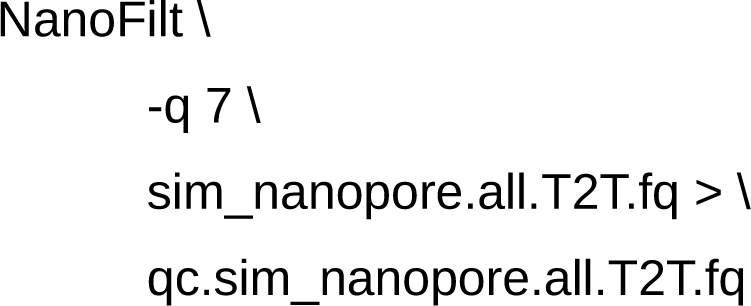

##### 4.1.2 Training of standard parameters

After filtering, more than 24,000 sequences remained, of which roughly 8% originated from shuffled reads. Before optimal alignment parameters were sought using LAST-TRAIN version 1256 (Supplementary Figure S2), using SeqKit, filtered reads were converted from a FASTQ to a FASTA format. The FASTA file was then aligned to the T2T-CHM13v2 database with LAST version 1256 using the optimal parameters which we extracted from the output of LAST-TRAIN (-p) (Supplementary Figure S2). LAST-SPLIT version 1256 was used to remove spurious alignments from the raw alignment file (Supplementary Figure S2). In order to find the best settings for LAST-SPLIT, the raw alignment file was filtered by using values of 0.99, 0.95, 0.90, 0.85, and 0.80 for LAST-SPLIT’s -m parameter.

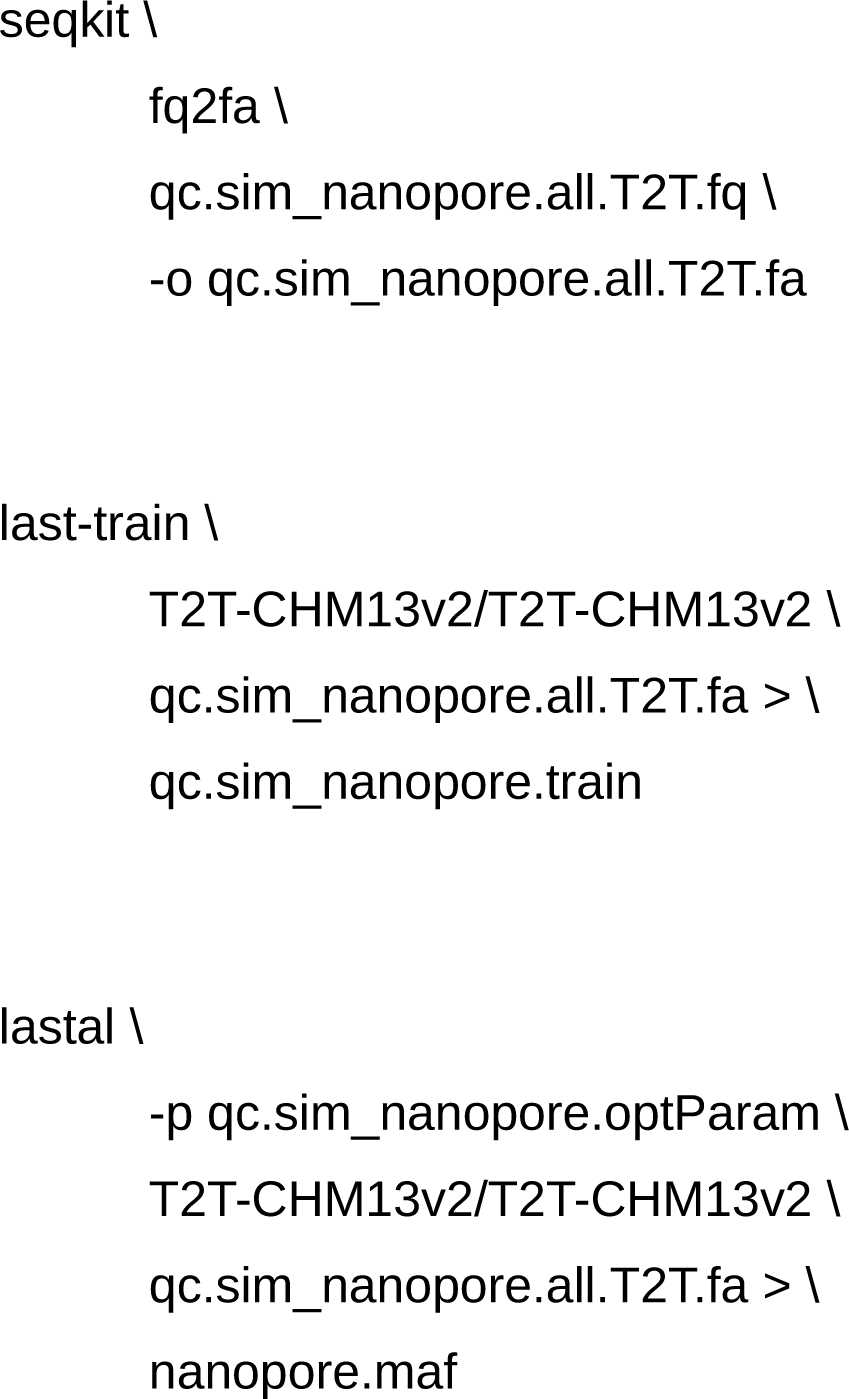

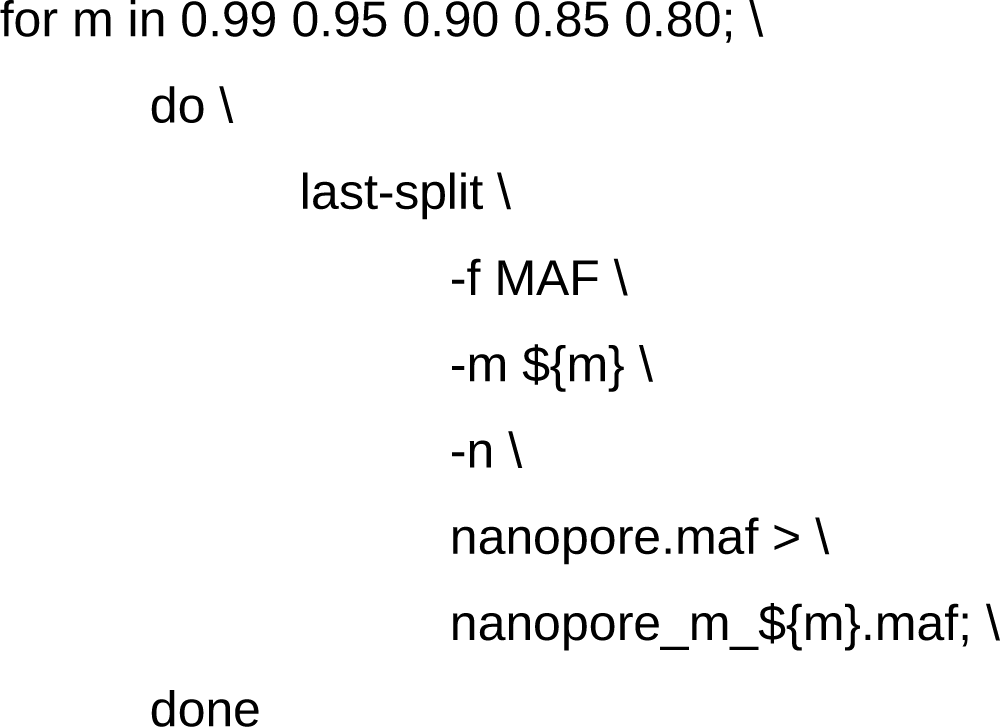

Aligned reads in the five filtered alignments and the raw alignment (-alignment) were extracted from the input FASTA file (-sequences) using MetaG’s filterReads.pl script (commit 49320ee; https://github.com/IOB-Muenster/MetaG/blob/master/metag_src/calculate/filterReads.pl; accessed 2024/01/22). The number of aligned human reads (more was better) and the number of aligned shuffled reads (less was better) were manually compared between all alignments (Supplementary Figure S2). The influence of the LAST-SPLIT settings in the tested range was minuscule. Nevertheless, the alignment filtered by the most permissive LAST-SPLIT setting (-m 0.99) was more than 200 times smaller than the native alignment (native alignment size: 168 GB). Smaller alignments meant faster processing time, so we included -m 0.99 together with the optimal alignment settings produced by LAST-TRAIN into our nanopore standard (commit 7ee14e9; https://github.com/IOB-Muenster/MetaG/blob/master/metag_src/install/files/ standard.config.filterHumanWGS.nanopore; accessed 2024/01/22).

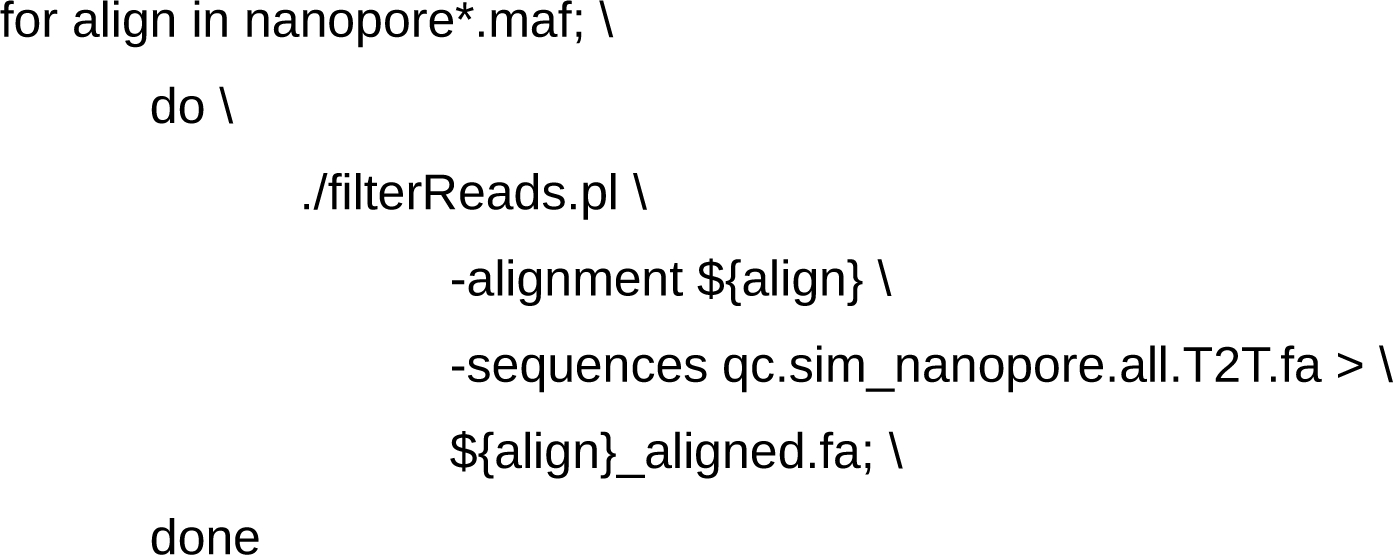

#### 4.2 Taxonomic classification

We downloaded the unaligned sequence FASTA files for bacteria, archaea, and fungi from RDP version 11.5 (https://rdp.cme.msu.edu/misc/resources.jsp; accessed 2021/11/24), the latest version of the database. We extracted and concatenated the files and replaced any semicolons with commas. The FASTA file (-input) was then processed with makeRDP.pl (commit 17d14ea; https://github.com/IOB-Muenster/MetaG/blob/master/supplemental/scripts/db/makeRDP.pl; accessed 2024/01/22) to build a database for MetaG.

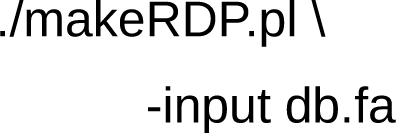

The script created a taxonomy file (tax.rdp.txt) and a reformatted sequence file (rdp.fa), which was subsequently processed with LASTdb (Supplementary Methods 4.1). The taxonomy was automatically modified to achieve a better taxonomic resolution. Namely, a separation of species and strain names was attempted (Manske, Grundmann, and Makałowski 2020). Furthermore, we aimed to use common expressions for non-informative taxa. For example, RDP provided several terms for an uncultured bacterial species from the genus *Acidimicrobium*: “uncultured bacterium,” “uncultured *Acidimicrobium* sp.,” and “uncultured prokaryote.” All these terms did not provide any relevant information for the taxonomy. So in these cases, we automatically set the species name to “uncultured.” Missing taxa in the lineage (domain, phylum, class, subclass, order, suborder, family, genus, species, or strain) and certain uninformative taxa were assigned to the special taxon “0.”

In order to predict pathogens (Manske, Grundmann, and Makałowski 2020), MetaG used the genome_lineage and genome_metadata files (ftp://ftp.bvbrc.org/ RELEASE_NOTES/; accessed 2023/10/30) from the Bacterial and Viral Bioinformatics Resource Center (BV-BRC) (Olson et al. 2023). Both files were processed with makeBVBRC.pl (commit: 275c4a9; https://github.com/IOB-Muenster/MetaG/blob/master/supplemental/scripts/db/makeBVBRC.pl; accessed 2024/01/22) to produce the pathogen taxonomy patho.BVBRC.txt, which only included pathogens from human hosts. In the taxonomy, the script automatically attempted to separate species and strain and to reformat non-informative taxa.

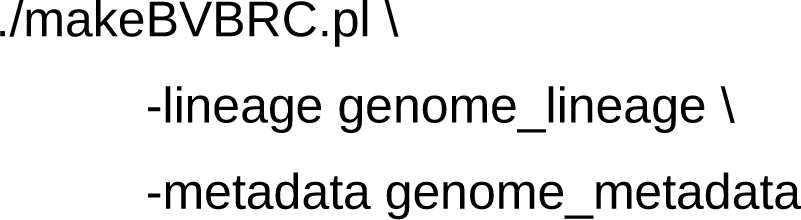

The pathogen database did not include any sequences and thus did not require any further preparations, including parameter tuning, before use with MetaG (-pdbPath). We ran MetaG using the RDP database (-ldb; -ltax) and a set of standard parameters (--config) on the PALMA II cluster in order to classify the filtered FASTA files (Supplementary Methods 4.1; Supplementary Figure S1). All filtered FASTA files were located in a single directory (-q) and processed simultaneously with the recursive mode of MetaG (--recursive). Training of standard parameters is described in the following sections (Supplementary Methods 4.2.1 and 4.2.2; Supplementary Figure S3).

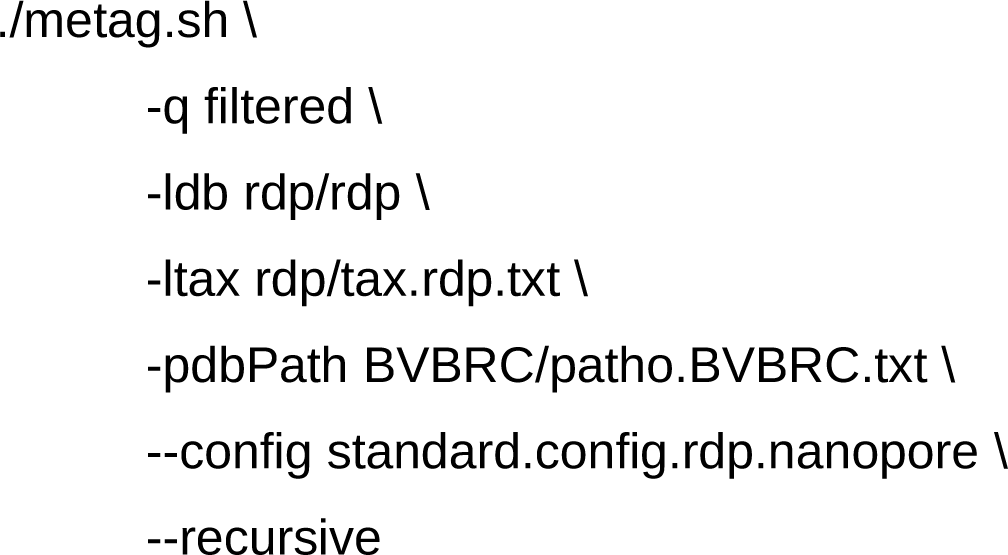

**Figure S3:**
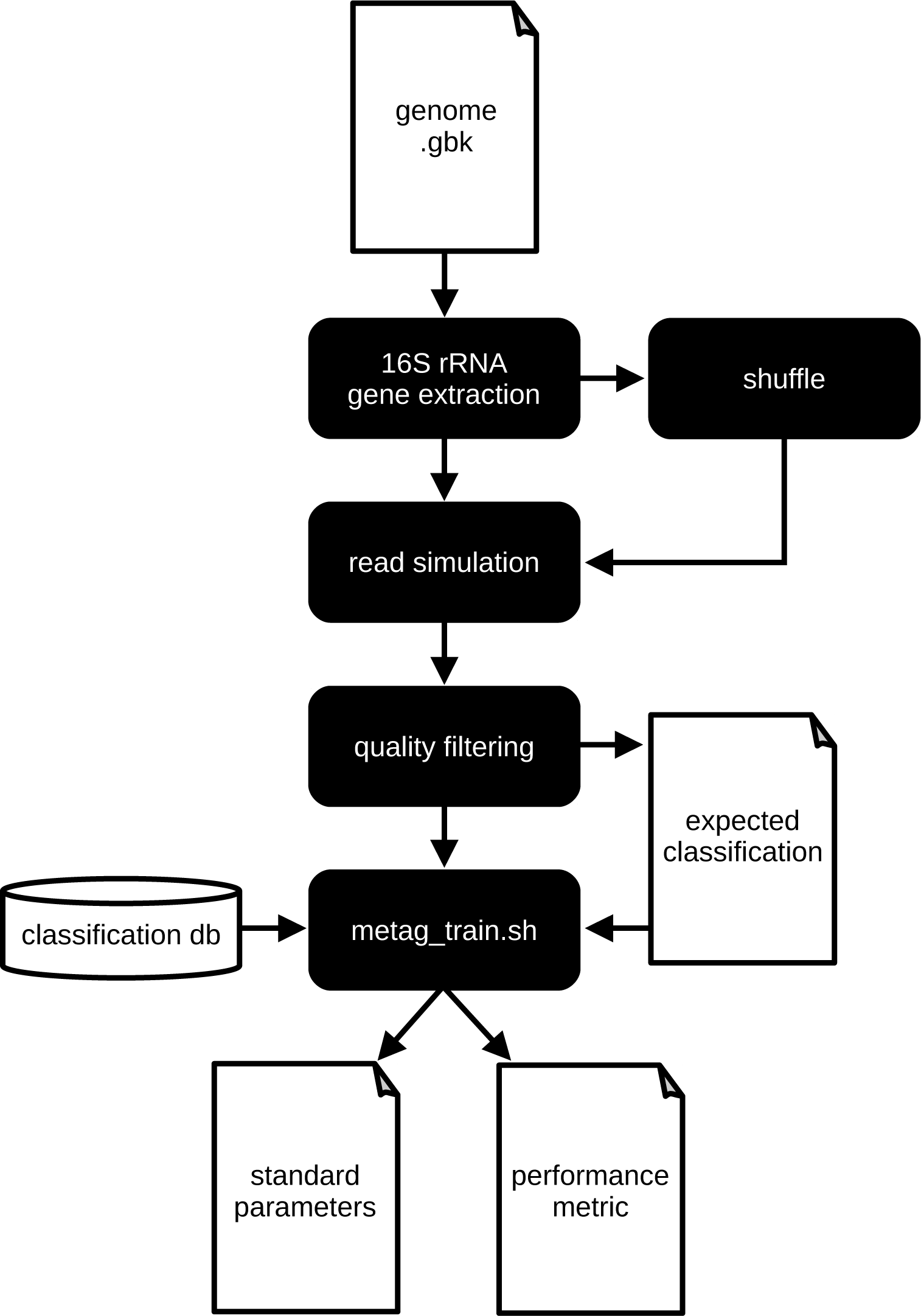
Workflow to generate the standard parameters for the taxonomic classifications. 16S ribosomal ribonucleic acid (rRNA) gene sequences were extracted from the genome “.gbk” files obtained from ATCC (https://genomes.atcc.org/; accessed 2022/01/04) and shuffled copies were generated. Sequencing of all copies was simulated and low-quality reads were filtered (Supplementary Methods 4.2.1). The retained reads, a file containing the expected classifications per read, and the classification database (“classification db”) were used to obtain standard parameters using metag_train.sh (commit: 9505087; https://github.com/IOB-Muenster/MetaG/blob/master/supplemental/scripts/train/metag-train.sh; accessed 2024/01/22). The performance metrics across multiple runs were compared in order to identify the overall best standard parameters (Supplementary Methods 4.2.2). This figure was generated with LibreOffice Draw 7.0.4.2 (https://www.libreoffice.org/discover/draw/; accessed 2024/03/22).

##### 4.2.1 *In silico* simulation of a mock sample

In order to identify the ideal parameters for taxonomic classifications with MetaG and the RDP database, we needed a collection of reads with known classifications. At the same time, we aimed to find generally applicable parameters that were neither specific to the current study nor to gut microbiota in general. Thus, we chose to use the 20 Strain Even Mix Genomic Material (Cat. No.: MSA-1002, ATCC, Manassas, USA) NGS standard. For each of the 20 strains, we obtained the reference annotations in GenBank (Sayers et al. 2023) flat file format from the manufacturer’s web portal (https://genomes.atcc.org/; accessed 2022/01/04). An ATCC user account was needed to obtain the data. From each flat file, we obtained the reference sequences for all 16S rRNA gene copies (Supplementary Figure S3) using parse-genbank_ATCC_sim.py (commit: 46d0677; https://github.com/IOB-Muenster/MetaG/blob/master/supplemental/scripts/train/parse-genbank_ATCC_sim.py; accessed 2024/06/07), which was a modified version of parse- genbank.py (original commit: 1142c6e; https://github.com/adina/tutorial-ngs-2014/blob/master/ncbi/parse-genbank.py; accessed 2022/01/03). The resulting FASTA files were split by each rRNA gene copy (-s1) using SeqKit. The provided commands illustrate the process for the example of a hypothetical genome “genome1”.

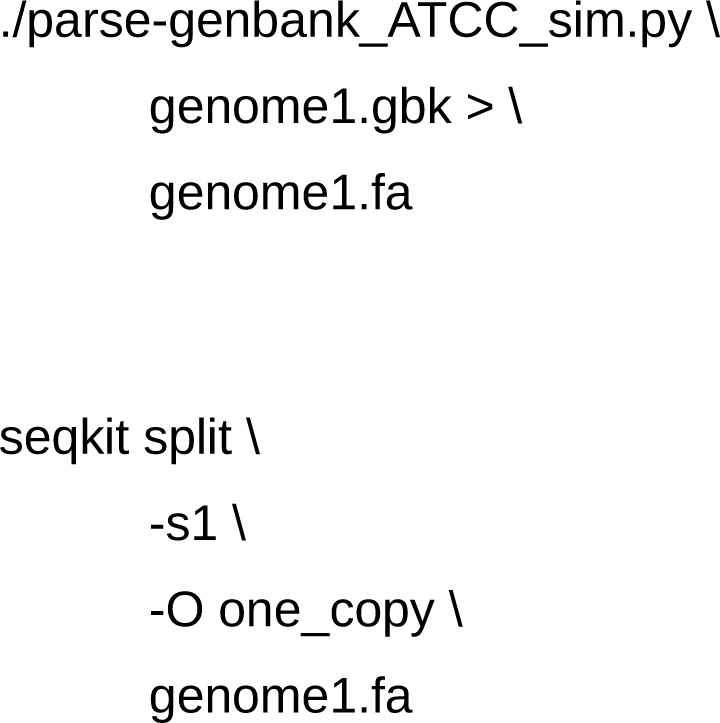

Additionally, a shuffled version of each split file was created (Supplementary Figure S3) using easel from HMMER (Supplementary Methods 4.1.1) in order to simulate sequences with no valid classifications. In reality, these could be artifacts or sequences from novel species which were not yet present in the reference database. Using Badread, we simulated nanopore sequencing (Supplementary Figure S3) at a target depth of 1,000x for each of the unshuffled rRNA gene FASTA files and at 100x for each of the shuffled rRNA gene FASTA files (--quantity). Badread did not have an amplicon simulation mode, so we set the read length (--length) to a very high number in order to force Badread to start the simulation at the beginning of the reference sequence (as exemplified below for an rRNA gene copy of a hypothetical genome “genome1”). We pooled all simulated nanopore reads and polished the headers using SeqKit.

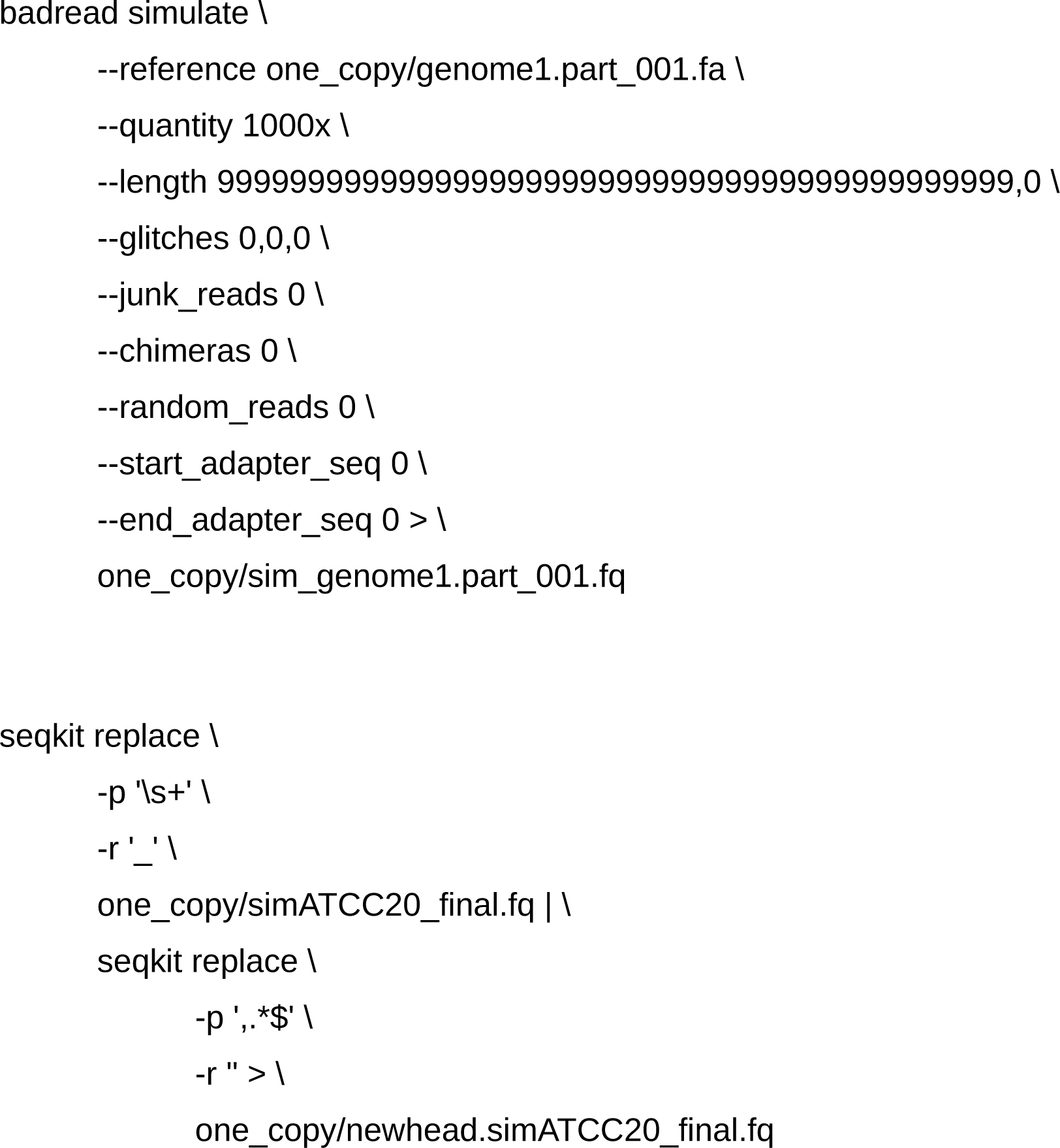

Reads were subject to the same quality filtering pipeline as described for the human filter database (Supplementary Methods 4.1.1; Supplementary Figure S3). The filtered sample contained more than 131,000 reads, of which roughly 9% originated from shuffled sequences.

##### 4.2.2 Automated training of standard parameters

The training script metag-train.sh (commit: 9505087; https://github.com/IOB-Muenster/ MetaG/blob/master/supplemental/scripts/train/metag-train.sh; accessed 2024/01/22) used the quality-filtered FASTQ file and a file containing the expected classifications to obtain the optimal MetaG parameters for nanopore metabarcoding analyses using the RDP database (Supplementary Figure S3). As reads were simulated, an expectation file was created that mapped each read to its expected classification. Shuffled reads were expected to be unclassified. The format of the expectation file was as follows: the first line started with “@@ranks” followed by a tab and a string identifying the ranks of the expected taxonomy: “domain;phylum;class;order;family;genus;species.” Subsequently, patterns that uniquely identified shuffled reads were listed on separate lines after a line only containing “@@negative.” Here, we simply listed the word “shuffled,” as it was present in the headers of all reads simulated from shuffled rRNA gene sequences. After a line only containing “@@positive,” expected classifications for classifiable reads were listed on separate lines. Each line contained a unique pattern followed by a tab and the expected classifications over all ranks (see “@@ranks”) separated by semicolon. Our script also supported the use of expected numbers of reads for the classifications, instead of unique patterns, in cases where the origin of each read was unknown. In our study, a unique pattern was based on the part of the sequence ID that was shared by all rRNA gene copies of an individual strain. To obtain each lineage, we manually extracted the taxonomy ID from the first entry in the history of cross references on the ATCC website for the particular strain (https://www.atcc.org/; accessed 2022/01/25) (Supplementary Table S2). We then used TaxonKit (Shen and Ren 2021) v0.8.0 to generate a lineage based on the National Center for Biotechnology Information (NCBI) taxonomy (Sayers et al. 2024) (https://ftp.ncbi.nih.gov/pub/taxonomy/; downloaded 2022/01/19; --data-dir) for each taxonomy ID. This is exemplified in the following command line for *Acinetobacter baumannii* ATCC 17978 (taxonomy ID: 400667).

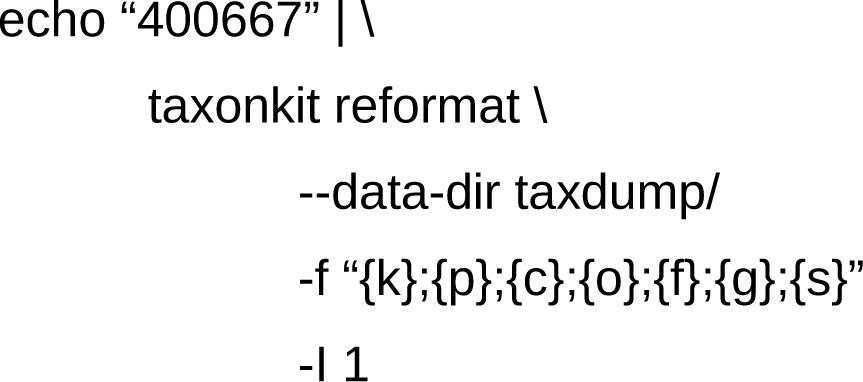

**Table S2:** ATCC strain name and NCBI taxonomy ID. The NCBI taxonomy (Sayers et al. 2024) ID for each of the strains in the 20 Strain Even Mix Genomic Material (Cat. No.: MSA-1002, ATCC, Manassas, USA) NGS standard was extracted from the history of cross references on www.atcc.org (2022/01/25).

| Strain | NCBI taxonomy ID |
| --- | --- |
| *Acinetobacter baumannii* ATCC 17978 | 400667 |
| *Bacillus pacificus* ATCC 10987 | 222523 |
| *Bacteroides vulgatus* ATCC 8482 | 435590 |
| *Bifidobacterium adolescentis* ATCC 15703 | 367928 |
| *Clostridium beijerinckii* ATCC 35702 | 864803 |
| *Cutibacterium acnes* ATCC 11828 | 1091045 |
| *Deinococcus radiodurans* ATCC BAA-816 | 243230 |
| *Enterococcus faecalis* ATCC 47077 | 474186 |
| *Escherichia coli* ATCC 700926 | 511145 |
| *Helicobacter pylori* ATCC 700392 | 85962 |
| *Lactobacillus gasseri* ATCC 33323 | 324831 |
| *Neisseria meningitidis* ATCC BAA-335 | 122586 |
| *Porphyromonas gingivalis* ATCC 33277 | 431947 |
| *Pseudomonas aeruginosa* ATCC 9027 | 287 |
| *Rhodobacter sphaeroides* ATCC 17029 | 349101 |
| *Schaalia odontolytica* ATCC 17982 | 411466 |
| *Staphylococcus aureus* ATCC BAA-1556 | 451515 |
| *Staphylococcus epidermidis* ATCC 12228 | 176280 |
| *Streptococcus agalactiae* ATCC BAA-611 | 208435 |
| *Streptococcus mutans* ATCC 700610 | 210007 |

The prepared FASTQ (-q) and expectation files (-exp) were used for fully automatic training of MetaG standard parameters. Training consisted of two major steps conducted by metag-train.sh. In the first step, LAST alignment parameters were trained using LAST-TRAIN. The optimal alignment parameters were used to generate a temporary alignment with LAST, which could be optionally filtered by LAST-SPLIT. In the second step, MetaG used this alignment to classify the input reads with all different combinations of e-value (-e), alignment score (-ac), and confidence cutoff (-cc) in a range specified by the user. Calculations were parallelized using GNU Parallel (Tange 2018) 20190922. For each parameter combination, the helper script metag-train.py (commit: 7ee14e9; https://github.com/IOB-Muenster/MetaG/blob/master/supplemental/scripts/train/metag-train.py; accessed 2024/01/22) calculated a performance metric based on the observed vs. expected classifications at a user-specified rank. The optimal parameter combination was defined as the strictest parameter combination (i.e. the highest values for -e, -ac, and -cc) with the highest performance metric and was written to an output configuration file together with the LAST parameters (Supplementary Figure S3). Two metrics were available for the training: for expectations where the origin of each read was known, the R_k_, a multiclass generalization of the Matthews correlation coefficient (Matthews 1975), (Gorodkin 2004) was used and for expectations where the origin of each read was unknown and expectations were purely based on read counts, the Bray-Curtis similarity index (BC) using the equation from H. Steinhaus as described by Motyka (Motyka 1947, 58) was used. Since the expected classifications for each read were known, we used the R_k_ in this study (-metric). We set the e-value range from e^-40^ to e^10^ (-e) with a step size of e^1^, the alignment score cutoff range from 0 to 1 (-ac) with a step size of 0.1, and the confidence cutoff range from 0 to 1 (-cc) with a step size of 0.1. On the PALMA II cluster, we performed training on the raw unfiltered LAST alignment (the command line is exemplified below) and on LAST alignments filtered by LAST-SPLIT (--lsplit 0.99, 0.95, 0.90, 0.85, and 0.80) considering only classifications at genus level (-rank). The maximum values for R_k_ (Supplementary Figure S3) were manually compared between the runs.

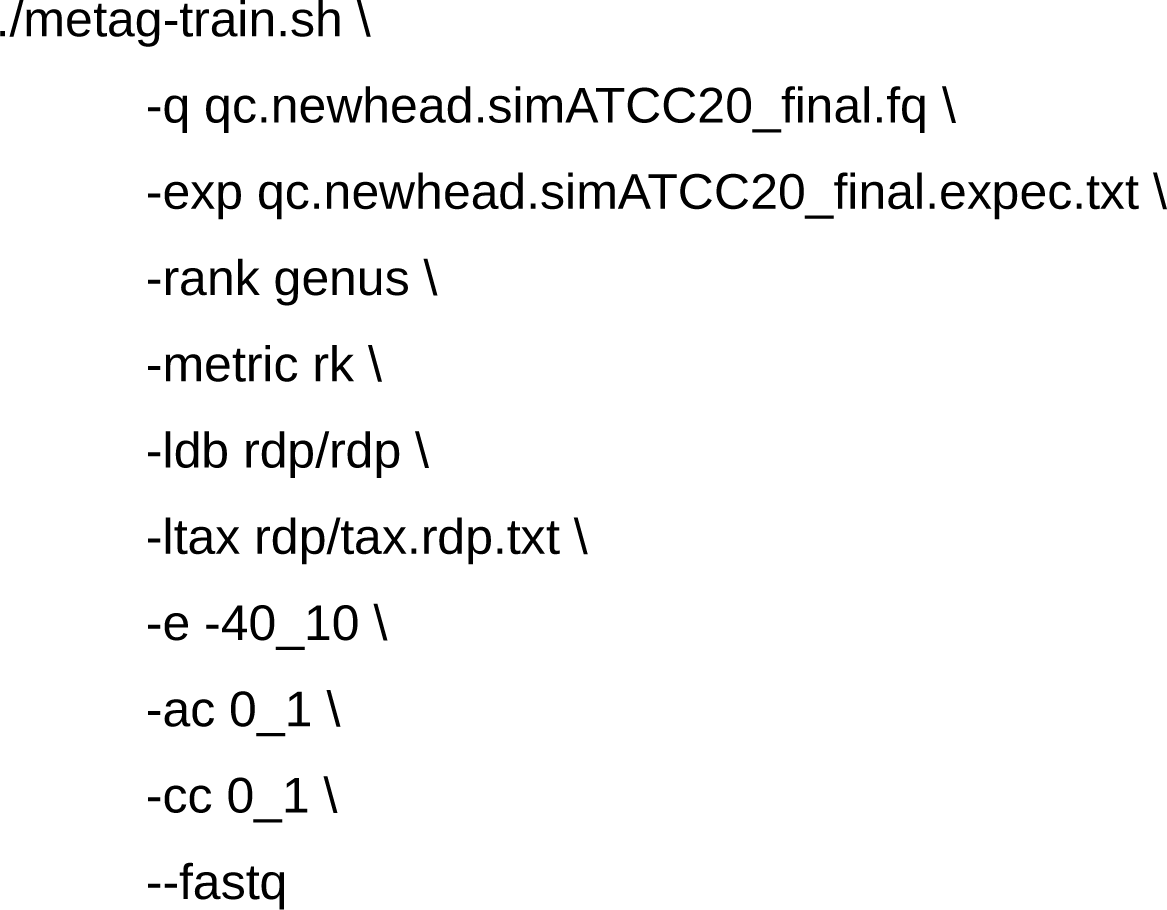

We found that performance differences between the runs were negligible and thus chose to include LAST-SPLIT filtering into the final standard parameters (commit: 7ee14e9; https://github.com/IOB-Muenster/MetaG/blob/master/metag_src/install/files/standard.config.rdp.nanopore; accessed 2024/01/22) in order to reduce alignment file size and thus reducing processing time, as discussed for the filter database (Supplementary Methods 4.1.2).

### 5 MetagenomicsDB - a central data management system

In order to store, explore, and export the data produced by our study, we created a central data management system called MetagenomicsDB (Figure 1): an import unit parsed the input files. After performing quality control, the data were inserted into a relational database which was used as the storage component. According to the user selection, (a subset of) stored records were exported by the respective unit in a format suitable for downstream statistical analyses. MetagenomicsDB could be operated using a web-based graphical interface (https://bioinformatics.uni-muenster.de/tools/ metagenomicsDB; accessed 2024/01/22) (Figure 1) which made our data management system accessible to all users, regardless of bioinformatics expertise. Our pipeline has already been described in the main part of this manuscript (Section 3). Here, we focus on the technical details which are necessary to understand in order to use MetagenomicsDB as a template for future microbiome studies.

#### 5.1 Data storage in a relational database

We stored the “pass” reads (Supplementary Methods Section 3.3), the classifications (Supplementary Methods 4.2; Supplementary Figure S1), the measurements (Supplementary Methods 2), and the patients’ and mothers’ metadata (Supplementary Methods 2) in a relational database. The database was implemented in PostgreSQL 14beta2 (https://www.postgresql.org/; accessed 2024/03/17) and its schema consisted of ten relations (Figure 2) and five materialized views (commit: 68b05c6; https://github.com/IOB-Muenster/MetagenomicsDB/blob/main/www-intern/db/schema.sql; accessed 2024/06/07). The following descriptions of the relations and views in the database can be reproduced by referring to the aforementioned schema.

##### 5.1.1 Relations in the database

In the following sections, names of relations, functions, and views will be italicized in order to distinguish them from regular English words. The *change* relation connected every record in the database with a username, a timestamp (column: “ts”), and an IP address (column: “ip”) (Figure 2), which we used for internal tracking of changes. The results from examining the patients and their mothers (Supplementary Methods 2) were stored in the *measurement* relation, which was connected via *sample* (many-to-one) to the patient metadata in *patient* (many-to-one; Figure 2). Patients could be uniquely identified by their patient ID, hospital code, and birthdate (columns: “alias,” “accession,” and “birthdate,” respectively) (Supplementary Table S1). Nevertheless, we later found that patients in our study could also be uniquely identified by using their patient ID, so we relied on the uniqueness of the patient ID in other parts of our implementation.

The *sample* relation contained the sample collection date (column: “createdate”), the examiner (column: “createdby”), and information if the sample was a control sample (column: “iscontrol”; Supplementary Methods 3.3). “iscontrol,” “createdate,” and the foreign key referencing the *patient* relation (column: “id_patient”) uniquely identified each sample. Since water-control samples only controlled for errors during sequencing and library preparation, by definition they had no measurements. Measurements that could change over the observation period were called variable measurements. Others remained constant and were thus termed static measurements. Both are shown in Supplementary Table S3.

**Table S3:** Static and variable measurements in MetagenomicsDB. Static and variable measurements stored in the *measurement* relation of MetagenomicsDB. The values for variable measurements could change over the observation period, while the values of static measurements remained constant. See Supplementary Table S1 for definitions.

| Measurement name | Type |
| --- | --- |
| Birth mode | Static |
| Maternal antibiotics during pregnancy | Static |
| Maternal body mass at delivery | Static |
| Maternal body mass before pregnancy | Static |
| Maternal illness during pregnancy | Static |
| Mother's birth date | Static |
| Mother's height | Static |
| Pregnancy order | Static |
| Sex | Static |
| Antibiotics | Variable |
| Body mass | Variable |
| Feeding mode | Variable |
| Probiotics | Variable |

Derived measurements (Supplementary Table S4) could be directly calculated from other measurement values. To reduce redundancy, static measurements were only present for the first sample of each patient (“meconium” sample) and derived measurements were only calculated in the views (Supplementary Methods 5.1.2) and not stored in *measurement*. In order to create a flexible database that could be easily adjusted for projects with different measurements, we stored static and variable measurements of a sample as rows in the *measurement* relation. This had the benefit of each sample being assigned to any number of measurements without modifying the database schema. Measurement names were normalized by a *type* relation that was linked to *measurement* (one-to-many) (Figure 2). *type* contained the measurement name (had to be unique), its type (“b” for boolean, “i” for integer, “s” for string, and “d” for date), and an array of options for menus in an internal admin web interface (column: “selection”). Measurements were uniquely identified by their foreign keys referencing *type* and *sample* (columns: “id_type” and “id_sample”, respectively).

**Table S4:** Derived measurements in MetagenomicsDB. The values of derived measurements stored in MetagenomicsDB were inferred from other measurement values. More information on the calculations is provided in Supplementary Methods 5.1.2.

| Derived measurement name |
| --- |
| Category of difference in body mass at delivery |
| Difference in body mass at delivery |
| Mother's age at delivery |
| Mother's pre-pregnancy BMI |
| Mother's pre-pregnancy BMI category |
| Z-score |
| Z-score category |
| Z-score subcategory |

Each entry in *sample* was connected to any number of sequences in the *sequence* relation (Figure 2). Apart from the nucleotides (column: “nucs”), base quality strings (column: “quality”), and metadata like run ID and barcode, it contained stored generated columns. These contained the average sequence error (column: “seqerr”) calculated via *f_calcseqerror* (Supplementary Methods 5.1.2) and the sequence length (column: “seqlen”). The values in stored generated columns were automatically recalculated on inserts and updates from other regular columns in the same row and relation (https://www.postgresql.org/docs/14/ddl-generated-columns.html; accessed 2024/01/22). Since sequence length and average sequence error directly depended on the nucleotide and quality strings in the same relation, respectively, generated columns reduced redundancy. Duplicates of run ID, barcode, read ID, and of the foreign key referencing *sample* (columns: “runid”, “barcode”, “readid”, and “id_sample”, respectively) were not allowed.

In its schema, MetagenomicsDB supported the taxonomic classification of a single sequence with multiple different programs (column: “program”) and/or databases (column: “database”) via the *classification* relation which was connected in a many-to-one relationship with *sequence* (Figure 2). The *classification* relation contained only one entry for each combination of program, database, and the foreign key referencing *sequence* (column: “id_sequence”). Taxa (column: “name”) and their respective ranks (column: “rank”) were stored in *taxonomy* (Figure 2). Entries in *taxonomy* were uniquely identified by columns called “name” and “rank.” In PostgreSQL 14beta2, null values were considered distinct in unique constraints (https://www.postgresql.org/docs/14/ddl-constraints.html#DDL-CONSTRAINTS-UNIQUE-CONSTRAINTS; accessed 2024/02/06), but we also wanted to catch duplicates of the same rank and of the null name. Thus, we used a unique index. Since *taxonomy* and *classification* had a many-to-many relationship, both relations were connected with the associative table *taxclass* (Figure 2). Isolated from the rest but *change* was the *standard* relation (Figure 2). It contained raw values from the WHO’s weight-for-age growth standards (Supplementary Methods 5.3.1), which were used to calculate the weight-for-age z-scores in the *v_measurements* view (Supplementary Methods 5.1.2). The unique constraint on *standard* comprised the columns “name” (name of the standard: “weight_for_age”), “sex,” and “age.”

##### 5.1.2 Views in the database

Five materialized views were needed for the functionalities of the web interface (Supplementary Methods 5.2). Materialization was chosen in order to improve the performance of database queries. The *v_lineages* view assembled a lineage from domain to strain for each sequence. Classifications per sample were supplemented with supporting read counts for each analysis workflow (analysis program and database). *v_samples* mainly joined the *patient* and *sample* relations, counted the number of sequencing reads per sample, and translated the sample collection dates to strings describing the sampling time point (“timepoint”). Time points were labeled relative to the birthdate of the respective patient in order to allow for convenient comparisons across patients. Definitions were fuzzy in order to account for minor variation in the sample collection date, but did not overlap. The following time points were defined (days after birth): “meconium” (0-1), “3d” (2-4), “2w” (10-18), “6w” (35-49), “3m” (79-101), “6m” (169-191), “9m” (259-281), and “1y” (354-376). Due to the purposefully fuzzy definitions, it was possible that multiple samples from the same patient and of the same sample type (case or control) were assigned to the same time point, without violating the unique constraint on *sample* (Supplementary Methods 5.1.1). This was not intended, since we expected a one-to-one mapping between sample collection dates for a patient and sampling time points for a given sample type. Thus, the view also contained a special column “isok.” The value in this column for the individual sample was false when no time point string could be assigned or when the aforementioned duplicate time point occurred. This column was used during insert and update operations on *sample* in order to automatically assess the validity of new or updated records.

*v_taxa* merged *patient*, *v_samples*, *sequence*, *classification*, *taxclass*, and *taxonomy* and was the main data source for the display of taxonomic classification results in the web interface (Supplementary Methods 5.2). It provided the results of the taxonomic classification for each sample analyzed by a specific combination of analysis program and database. Classifications were split by rank. Supporting read counts for each taxon were supplemented with statistics on the reads: minimum, maximum, and average values were given for the read length and for the average read quality. The average read quality (readQ) of a read with length N was calculated from the Phred score (Ewing and Green 1998) of each base (q_i_) following the equation used by the Guppy basecaller (https://labs.epi2me.io/quality-scores/; accessed 2024/01/24):

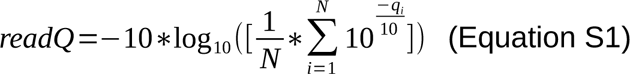

According to Supplementary Equation S1, the average read quality was the Phred score of the average sequence error. In MetagenomicsDB, the function *f_calcseqerror* first calculated the average sequence error of every entry in *sequence* (column “seqerr” in *sequence*; Supplementary Methods 5.1.1). The minimum and maximum average read qualities were defined in *v_taxa* as the Phred scores of the maximum and minimum average sequence error, respectively. To calculate the mean of readQ, average sequence errors from all supporting sequences were again averaged and expressed as a Phred score.

In order to query the *measurement* relation and to calculate the derived measurements (Supplementary Table S4), we created the *v_measurements* view. The view contained patient and sample metadata, as well as the following derived measurements: “mother’s age at delivery” (in years) was calculated based on the patient’s birthdate and the birthdate of the mother (“birth date” and “mother’s birth date,” Supplementary Table S1) assuming a year consisted of 365.25 days. Using the “maternal body mass before pregnancy” and “mother’s height” (Supplementary Table S1), we calculated the maternal pre-pregnancy body mass index (BMI) (“mother’s pre-pregnancy BMI”) as weight (in kg) divided by the height (in m) squared (Keys et al. 1972). Based on the ICD-11 (World Health Organization 2022, 348-349,1807), mothers were assigned to one of four groups according to their BMIs (“mother‘s pre-pregnancy BMI category”): “underweight” (BMI < 18.5 kg/m^2^), “normal weight” (18.5 kg/m^2^ <= BMI < 25.0 kg/m^2^), “overweight” (25.0 kg/m^2^ <= BMI < 30.0 kg/m^2^), and “obesity” (30.0 kg/m^2^ <= BMI). Additionally, the “difference in body mass at delivery” (Δm_del_) was calculated as the difference between the “maternal body mass at delivery” and the “maternal body mass before pregnancy” (Supplementary Table S1). In accordance with previously published guidelines (Institute of Medicine and National Research Council 2009, tbls. 7–3), the weight gain or loss during pregnancy was then assessed using the BMI („category of difference in body mass at delivery“). Mothers carrying multiple fetuses were not present in the study cohort (Supplementary Methods 1):

- “not enough”

◦ BMI < 18.5 kg/m^2^ and Δm_del_ < 12.5 kg
◦ 18.5 kg/m^2^ <= BMI < 25.0 kg/m^2^ and Δm_del_ < 11.5 kg
◦ 25.0 kg/m^2^ <= BMI < 30.0 kg/m^2^ and Δm_del_ < 7.0 kg
◦ 30 kg/m^2^ <= BMI and Δm_del_ < 5.0 kg)
- “appropriate”

◦ BMI < 18.5 kg/m^2^ and 12.5 kg <= Δm_del_ <= 18.0 kg
◦ 18.5 kg/m^2^ <= BMI < 25.0 kg/m^2^ and 11.5 kg <= Δm_del_ <= 16.0 kg
◦ 25.0 kg/m^2^ <= BMI < 30.0 kg/m^2^ and 7.0 kg <= Δm_del_ <= 11.5 kg
◦ 30 kg/m^2^ <= BMI and 5.0 kg <= Δm_del_ <= 9.0 kg)
- “too much”

◦ BMI < 18.5 kg/m^2^ and Δm_del_ > 18.0 kg
◦ 18.5 kg/m^2^ <= BMI < 25.0 kg/m^2^ and Δm_del_ > 16.0 kg
◦ 25.0 kg/m^2^ <= BMI < 30.0 kg/m^2^ and Δm_del_ > 11.5 kg
◦ 30 kg/m^2^ <= BMI and Δm_del_ > 9.0 kg

In order to divide the patients into categories with appropriate and deficient growth, the weight-for-age z-score was calculated by the function *f_calczscore*. It was implemented based on the equations provided by the WHO (WHO Multicentre Growth Reference Study Group 2006, 302–3). This function required the patient’s body mass (in kg) at a specific time point (“body mass”, Supplementary Table S1), and lambda, mu, and sigma values generated by the WHO (Supplementary Methods 5.3.1). Lambda (column: “l”), mu (column: “m”), and sigma (column: “s”) were queried from the *standard* relation based on the sex and the exact age of the patient in days (Figure 2). Patients for which the weight-for-age z-score was not a suitable measure (e.g. those with oedema (World Health Organization and United Nations Children’s Fund (UNICEF) 2019, tbl. 6)) were not present in the database (Supplementary Methods 1). The “z-score category” was defined according to the criteria provided in Supplementary Methods 1. SGA children were analyzed in greater depth to see if (and when) they caught up in growth with AGA children. To achieve this, we created sub-categories for SGA children (“z-score subcategory”) based on the z-score, its change over time, and the sampling time point. By definition, the earliest time point at which catch-up could occur was in the second sample (“3d”) and “SGA” was used as the sub-category name for the “meconium” sample. At catch-up, patients needed to have a z-score of more than negative two. Furthermore, the z-score difference between the lowest z-score at a previous measurement and the current z-score had to be at least 0.67. If the catch-up event happened in the first six month, it was termed “early catch-up,” otherwise it was called “late catch-up.” Before the catch-up event, but only from the “3d” sample on, SGA children were assigned to the “no catch-up” sub-category. The same applied to children who never showed catch-up growth during our study.

*v_measurements* followed the aforementioned definition that static measurements were only assigned to the “meconium” sample (Supplementary Methods 5.1.1; Supplementary Table S3). Since all derived measurements (Supplementary Table S4) were based directly or indirectly on statics, all but the z-score and its derivatives were only present for the first sample. In contrast, *v_metadata* assigned a subset of these measurements to all sampling time points of a patient. Namely, this included “sex,” “birth mode,” “mother’s age at delivery,” “mother’s pre-pregnancy BMI category,” “category of difference in body mass at delivery,” “pregnancy order,” “maternal illness during pregnancy,” and “maternal antibiotics during pregnancy” (Supplementary Tables S3 and S4). Furthermore, the view added new measurements containing the classification program, the classification database, and also the time point itself. The special format of the view was needed for the export to the MicrobiomeAnalyst (Lu et al. 2023) (Supplementary Methods 5.2). Otherwise, *v_metadata* largely displayed the same data as *v_measurements*. Both views did not contain control samples, as these had no measurements (Supplementary Methods 5.1.1).

#### 5.2 Web interface

The web interface and the export to the MicrobiomeAnalyst have been extensively described in the main part of this manuscript (Sections 3.3 and 3.4). Here we want to briefly outline some more technical aspects. The columns in all three tabs of the web interface are described in Supplementary Table S5. The patients tab of the web interface was based on information from the *patient* relation and *v_measurements* (Supplementary Methods 5.1.1 and 5.1.2). *v_samples* and *v_measurements* (Supplementary Methods 5.1.2) were integral to the samples tab. The taxa tab was based on the views *v_taxa* and *v_lineages* (Supplementary Methods 5.1.2).

**Table S5:**
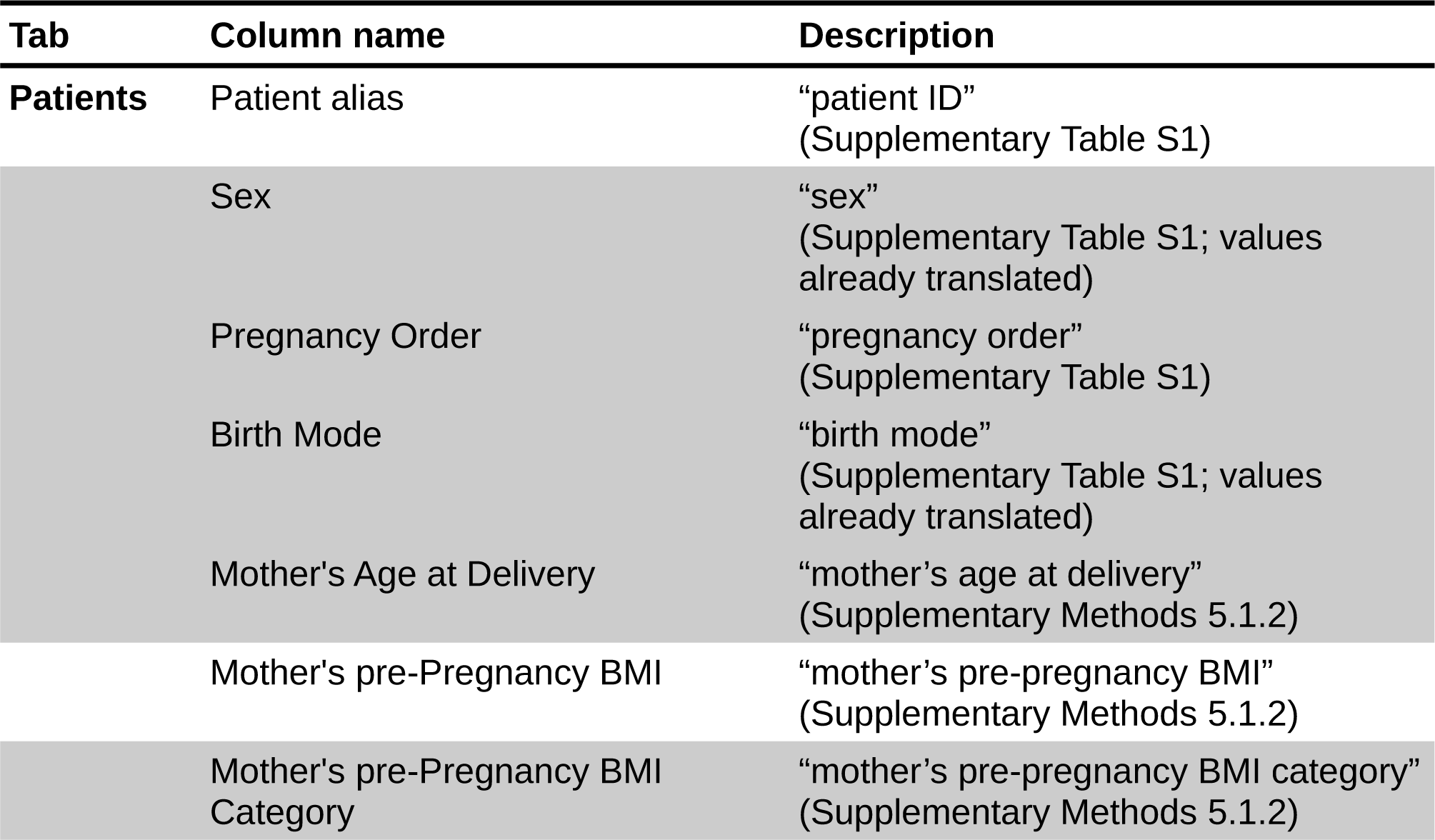
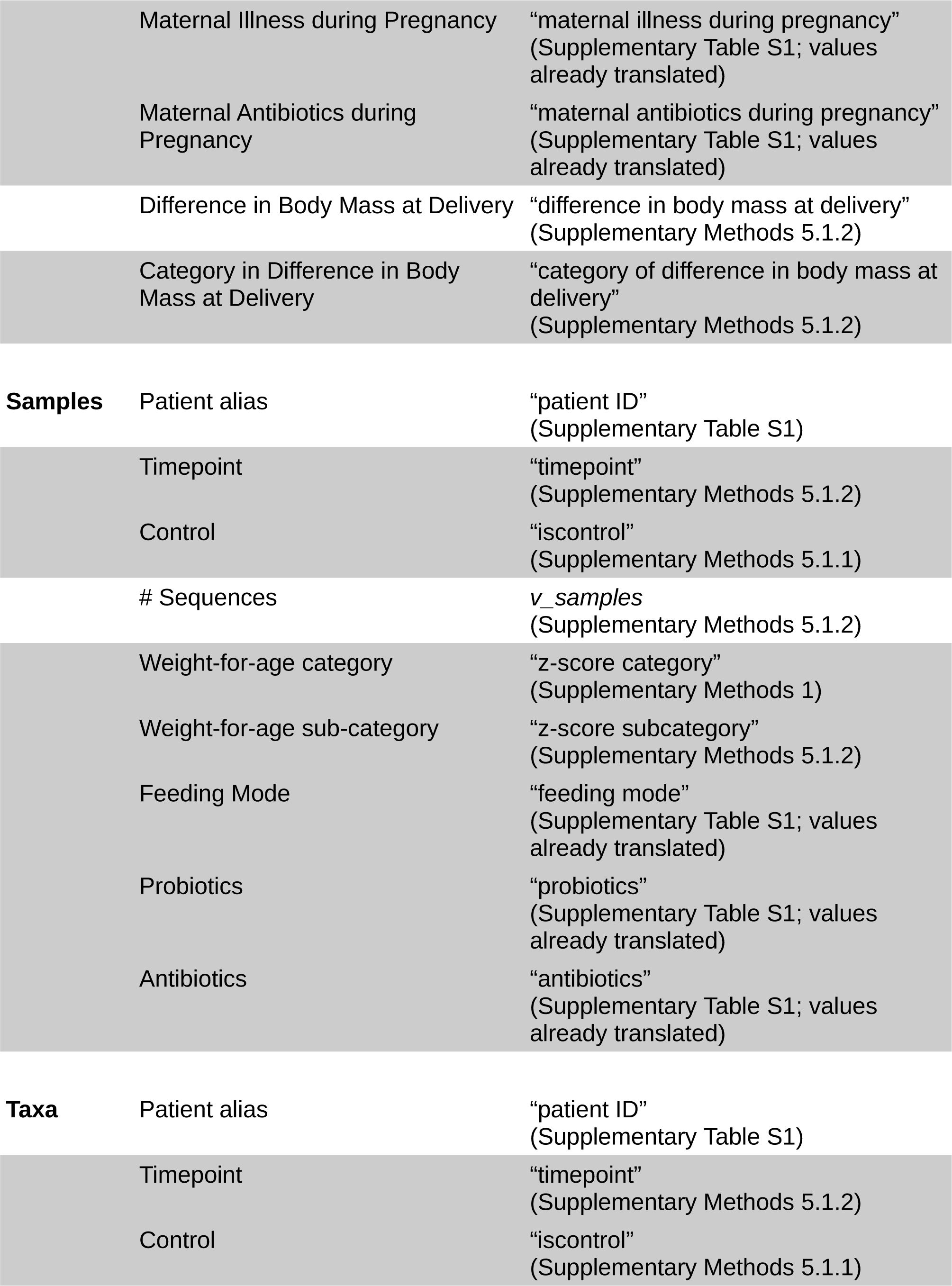
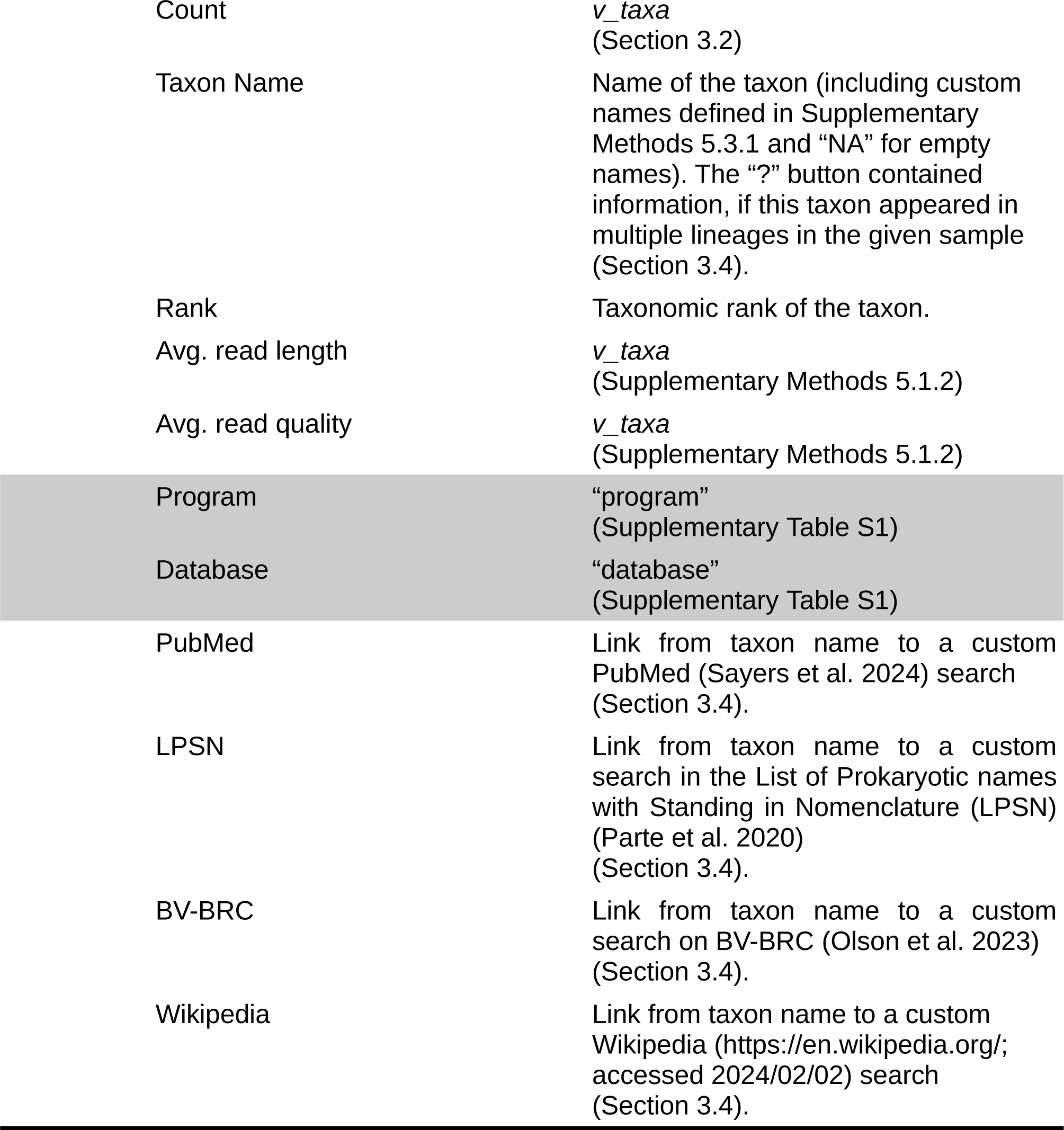
Columns in the tabs of the MetagenomicsDB web interface. Definitions of the columns in the tabs of the MetagenomicsDB web interface. Grey shading is used for metadata values exported to the MicrobiomeAnalyst (Lu et al. 2023).

| Tab | Column name | Description |
| --- | --- | --- |
| Patients | Patient alias | “patient ID”<br>(Supplementary Table S1) |
|  | Sex | “sex”<br>(Supplementary Table S1; values already translated) |
|  | Pregnancy Order | “pregnancy order”<br>(Supplementary Table S1) |
|  | Birth Mode | “birth mode”<br>(Supplementary Table S1; values already translated) |
|  | Mother’s Age at Delivery | “mother’s age at delivery”<br>(Supplementary Methods 5.1.2) |
|  | Mother’s pre-Pregnancy BMI | “mother’s pre-pregnancy BMI”<br>(Supplementary Methods 5.1.2) |
|  | Mother’s pre-Pregnancy BMI Category | “mother’s pre-pregnancy BMI category”<br>(Supplementary Methods 5.1.2) |

**Table S5: Columns in the tabs of the MetagenomicsDB web interface (continued).**
|  |  |  |
| --- | --- | --- |
|  | Maternal Illness during Pregnancy | "maternal illness during pregnancy"<br>(Supplementary Table S1; values already translated) |
|  | Maternal Antibiotics during Pregnancy | "maternal antibiotics during pregnancy"<br>(Supplementary Table S1; values already translated) |
|  | Difference in Body Mass at Delivery | "difference in body mass at delivery"<br>(Supplementary Methods 5.1.2) |
|  | Category in Difference in Body Mass at Delivery | "category of difference in body mass at delivery"<br>(Supplementary Methods 5.1.2) |
| **Samples** | Patient alias | "patient ID"<br>(Supplementary Table S1) |
|  | Timepoint | "timepoint"<br>(Supplementary Methods 5.1.2) |
|  | Control | "iscontrol"<br>(Supplementary Methods 5.1.1) |
|  | # Sequences | *v_samples*<br>(Supplementary Methods 5.1.2) |
|  | Weight-for-age category | "z-score category"<br>(Supplementary Methods 1) |
|  | Weight-for-age sub-category | "z-score subcategory"<br>(Supplementary Methods 5.1.2) |
|  | Feeding Mode | "feeding mode"<br>(Supplementary Table S1; values already translated) |
|  | Probiotics | "probiotics"<br>(Supplementary Table S1; values already translated) |
|  | Antibiotics | "antibiotics"<br>(Supplementary Table S1; values already translated) |
| **Taxa** | Patient alias | "patient ID"<br>(Supplementary Table S1) |
|  | Timepoint | "timepoint"<br>(Supplementary Methods 5.1.2) |
|  | Control | "iscontrol"<br>(Supplementary Methods 5.1.1) |

**Table S5: Columns in the tabs of the MetagenomicsDB web interface (continued).**
|  |  |
| --- | --- |
| Count | *v_taxa*<br>(Section 3.2) |
| Taxon Name | Name of the taxon (including custom names defined in Supplementary Methods 5.3.1 and “NA” for empty names). The “?” button contained information, if this taxon appeared in multiple lineages in the given sample (Section 3.4). |
| Rank | Taxonomic rank of the taxon. |
| Avg. read length | *v_taxa*<br>(Supplementary Methods 5.1.2) |
| Avg. read quality | *v_taxa*<br>(Supplementary Methods 5.1.2) |
| Program | “program”<br>(Supplementary Table S1) |
| Database | “database”<br>(Supplementary Table S1) |
| PubMed | Link from taxon name to a custom PubMed (Sayers et al. 2024) search (Section 3.4). |
| LPSN | Link from taxon name to a custom search in the List of Prokaryotic names with Standing in Nomenclature (LPSN) (Parte et al. 2020) (Section 3.4). |
| BV-BRC | Link from taxon name to a custom search on BV-BRC (Olson et al. 2023) (Section 3.4). |
| Wikipedia | Link from taxon name to a custom Wikipedia (https://en.wikipedia.org/; accessed 2024/02/02) search (Section 3.4). |

From the patients and samples tab, data could be exported to MicrobiomeAnalyst. The export was based on the program exportSGA.pl (commit: 68b05c6; https://github.com/IOB-Muenster/MetagenomicsDB/blob/main/bin/exportSGA.pl; accessed: 2024/06/07). Given a set of sample IDs, it exported classifications and sample metadata from the views *v_lineages*, *v_samples*, and *v_metadata* (Supplementary Methods 5.1.2; Supplementary Table S5). By default, “FILTERED” taxa (Supplementary Methods 5.3.1) and internal control samples (Supplementary Methods 3.3) were not exported. This behavior could be adjusted according to the user’s preferences. When the blacklist was explicitly defined as empty (--blacklist ‘’), all taxa were exported. Control samples could be exported by adding --keepctrl to the command line of exportSGA.pl. The following example shows the export of a hypothetical sample with the database ID “1” to MicrobiomeAnalyst (--format) using default settings.

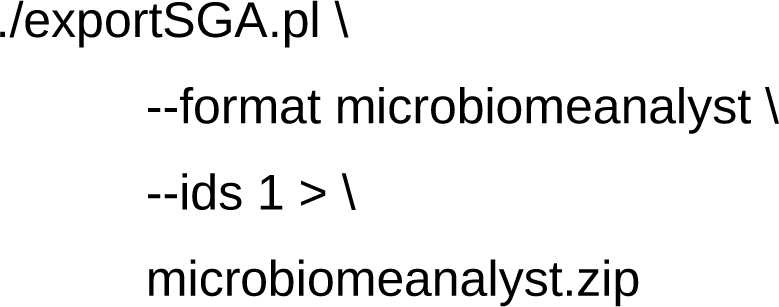

#### 5.3 Populating and modifying the database contents

In order to populate the database and to modify its contents, we developed the program importSGA.pl (commit: 68b05c6; https://github.com/IOB-Muenster/MetagenomicsDB/blob/ main/bin/importSGA.pl; accessed: 2024/06/07). It automated the quality control of the input data as much as possible in order to reduce human error. The command line was the same for all operations which aimed to synchronize data between the input files and the database, as our pipeline autonomously decided which operation (insert, update, or ignore) was necessary (Supplementary Methods 5.3.2). During our study, we did not require the deletion of records from the database. Thus, to avoid unintentional deletions, we did not implement this option into the program. In the following section, we will describe the input data and the command line parameters of importSGA.pl, while Supplementary Methods 5.3.2 will focus on the technical aspects of insert and update operations.

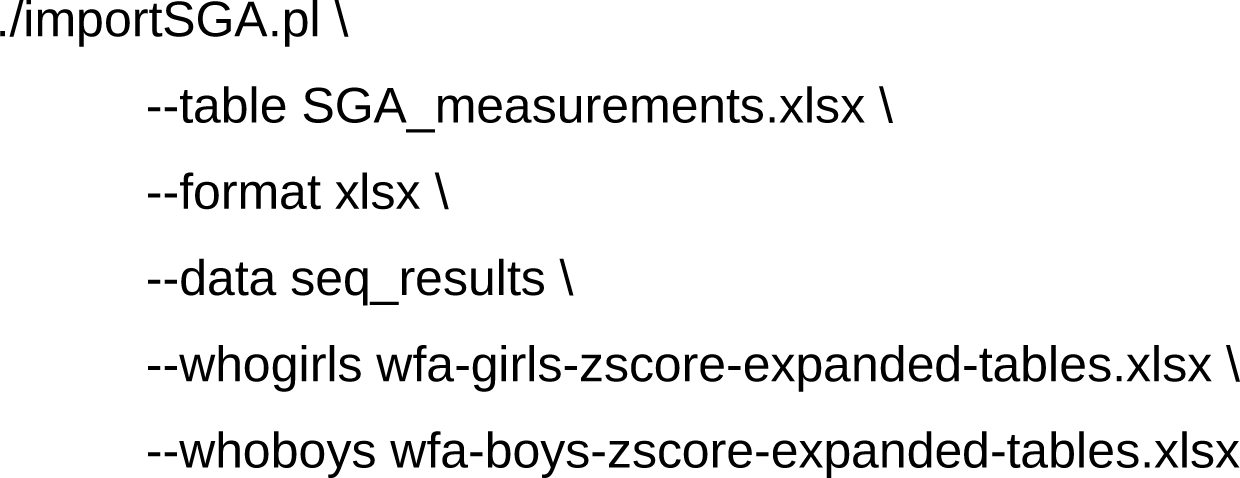

##### 5.3.1 Definitions and format of the input data

Weight-for-age growth standards from the WHO (Supplementary Table S6) were required to calculate the z-scores (Supplementary Methods 5.1.2). The files for boys (https://cdn.who.int/media/docs/default-source/child-growth/child-growth-standards/ indicators/weight-for-age/expanded-tables/wfa-boys-zscore-expanded-tables.xlsx; accessed 2024/01/30) and girls (https://cdn.who.int/media/docs/default-source/child-growth/child-growth-standards/indicators/weight-for-age/expanded-tables/wfa-girls-zscore-expanded-tables.xlsx; accessed 2024/01/30) were downloaded and provided to importSGA.pl via --whoboys and --whogirls, respectively. Both parameters were optional so the standards were only necessary during the initialization of the database from scratch or when the standards were supposed to be updated. Nevertheless, if present, both needed to be given. Age in days, lambda, mu, and sigma values were extracted from the files and stored for insertion into the *standard* relation together with the name of the standard (“weight_for_age”) and the sex (columns: “age,” “l,” “m,” “s,” “name,” and “sex,” respectively) (Figure 2).

**Table S6:** Permitted input file formats for importSGA.pl. Permitted file types for the input of importSGA.pl (commit: 68b05c6; https://github.com/IOB-Muenster/ MetagenomicsDB/blob/main/bin/importSGA.pl; accessed: 2024/06/07).

| Data type | File format | Compression | Nested archives |
| --- | --- | --- | --- |
| Weight-for-age growth standards | „.xlsx“ | - | - |
| Sequences | “.fastq“ | “.zip“, “.gz“, “.bz2“ | “.zip“, “.gz“, “.bz2“ |
| Classifications | calc.LIN.txt from MetaG | “.zip“, “.gz“, “.bz2“ | “.zip“, “.gz“, “.bz2“ |
| Measurements | “.xlsx“, “.xls“, “.ods“, “.sxc“, “.csv“ | “.zip“, “.gz“, “.bz2“ | - |

Unfiltered FASTQ “pass” files (Supplementary Methods 3.3; Supplementary Table S6) and the classifications of their filtered counterparts (Supplementary Methods 4.2; Supplementary Table S6) were located in a single directory, which could optionally be compressed to an archive (“.zip” or “.tar”) and was passed to importSGA.pl as --data. No nested archives were allowed. If the directory was compressed, the type of archive was inferred from the file extension or the archive’s magic number. The files inside of the directory had to be organized in subdirectories with unique names. In our case, names were based on the run and barcode of the samples (column: “number of run and barcode,” Supplementary Table S1), so that importSGA.pl could use this information to uniquely assign files to a sample (see below). The FASTQs and MetaG results from sequencing the control of each run (Supplementary Methods 3.3) were assigned to the special barcode 99, which was typically referenced by multiple control samples of the same run (Supplementary Methods 5.3.2). In contrast, sequences and classifications from the genuine stool samples could not be referenced by multiple samples. All FASTQ files inside a matching subdirectory were first extracted (Supplementary Table S6), if applicable, and then concatenated. To avoid the insertion of duplicate data, FASTQ files needed to have unique names, irrespective of the file extensions. We extracted read IDs, sequences, quality strings, and selected metadata from the sequence headers (“runid,” “barcode,” “flow_cell_id,” and “basecall_model_version_id”) for storage in the *sequence* relation (columns: “readid,” “nucs,” “quality,” “runid,” “barcode,” “flowcellid,” “callermodel,” respectively) (Figure 2).

We used the “calc.LIN.txt” files from MetaG which were based on the filtered FASTQ “pass” files (Supplementary Methods 4.2) to obtain the classifications of each read. Classification files had to be organized as described for the FASTQs and could also be compressed (Supplementary Table S6). We extracted the classifications for each read and the ranks domain, phylum, class, subclass, order, suborder, family, genus, species, and strain. For each of these ranks, reads that could not be at all classified and reads that could not be classified from the respective rank onward were assigned to the special taxon “UNMATCHED.” In two other cases, special taxon names were used. In the case the name of a taxon was empty (“0” in classification database (Supplementary Methods 4.2) which was encoded as ”unclassified” in MetaG output), it was assigned to an undefined value, which later became a null value in the database. If a read appeared in the FASTQ files, but not in the respective classification file, it had been removed by the MetaG filtering workflow (Supplementary Methods 4.1). In such a case, we set the taxon names over all ranks to “FILTERED.” If a read appeared in the classification file, but not in the associated FASTQ files, FASTQ and classification files did not match and an error appeared.

The measurement file (Supplementary Tables S1 and S6) provided as --table was essential for importSGA.pl. Not only did it contain patient and maternal metadata and information on the samples and measurements, but it also contained the directory patterns (column: “number of run and barcode,” Supplementary Table S1) which were used to identify the matching FASTQ and MetaG classification files for each sample (see above). The file format (Supplementary Table S6) was automatically guessed from the extension of the input file, but could also be provided using --format. While the organization of FASTQ and MetaG classification files could also be easily adapted by future projects, the format of the measurement file was expected to be project-specific. Thus in the following, this file and the general algorithm for parsing it will be described in more detail. The file contained a single sheet with a header in the first row. The parseTable function in the Sga.pm module (commit: 68b05c6; https://github.com/IOB-Muenster/MetagenomicsDB/blob/main/lib/perl/MetagDB/Sga.pm; accessed: 2024/06/07) listed columns which were supposed to be extracted from the table file. Thus, it was sensitive to changes of the format of the measurement file (e.g. due to deletions or additions of columns). Target columns were defined by their index in the input file and assigned to one of four groups. A single “id” column was expected to uniquely identify each patient and a single “timepoint” column contained the sample collection date. Rows with no value in the “id” column were skipped and a sample collection date had to be present in all rows. The rest of the columns were divided into “measurement” and “static,” depending on the fact if column values were supposed to change between samples of a patient or not (Supplementary Table S 7). Albeit different, this definition had some overlap with the definition of static and variable measurements presented earlier (Supplementary Table S3; Supplementary Methods 5.1.1). However, the definition in parseTable also comprised columns which were not stored in measurement (Supplementary Tables S3 and S7).

**Table S7:** Names, categories, and target relations of columns in the measurement table. Column names in the measurement table (Supplementary Table S1), their respective category in the parseTable function of Sga.pm (commit: 68b05c6; https://github.com/IOB-Muenster/MetagenomicsDB/blob/ main/lib/perl/**MetagDB**/Sga.pm; accessed: 2024/06/07), and the relation the value was stored in (if applicable). Column names with gray shading were not extracted from the input file.

| Column name | Category | Relation |
| --- | --- | --- |
| patient ID | id | *patient* |
| hospital code | static | *patient* |
| birth date | static | *patient* |
| sex | static | *measurement* |
| time point |  |  |
| sample collection date | timepoint | *sample* |
| body mass | measurement | *measurement* |
| body mass z-score |  |  |
| catch-up |  |  |
| birth mode | static | *measurement* |
| feeding mode | measurement | *measurement* |
| probiotics | measurement | *measurement* |
| antibiotics | measurement | *measurement* |
| mother's birth date | static | *measurement* |
| mother's age at delivery |  |  |
| maternal body mass before pregnancy | static | *measurement* |
| maternal body mass at delivery | static | *measurement* |
| mother's height | static | *measurement* |
| mother's pre-pregnancy BMI |  |  |
| pre-pregnancy BMI category |  |  |
| difference in body mass at delivery |  |  |
| category of difference in body mass at delivery |  |  |
| pregnancy order | static | *measurement* |
| maternal illness during pregnancy | static | *measurement* |
| maternal antibiotics during pregnancy | static | *measurement* |
| program | measurement | *classification* |
| database | measurement | *classification* |
| number of run and barcode | measurement |  |

Derived measurements (Supplementary Table S4) constituted the vast majority of columns which were not read from the table file (Supplementary Table S7), since they were later calculated within the database (Supplementary Methods 5.1.1 and 5.1.2). If columns other than the “timepoint” column contained dates, they could be specified by their name in order to verify the correct extraction from the input file. This was the case for “birth date” and “mother’s birth date.”

##### 5.3.2 Inserts and updates of database records

First, the database was created from its schema (Supplementary Methods 5.1). Host, user, database name, and password were stored in the “.pg_service.conf” file (https://www.postgresql.org/docs/14/libpq-pgservice.html; accessed 2024/01/30) under the service name “metagdb.” So that adjustments to different computing environments were straightforward, our script importSGA.pl used this PostgreSQL configuration file to connect to the database using its service name. Directly after the connection was established, an exclusive transaction-level advisory lock was acquired in order to prevent multiple instances of importSGA.pl to write to the database at the same time. The lock was automatically released at the end of the transaction by the database management system (https://www.postgresql.org/docs/14/explicit-locking.html; accessed 2024/02/01), even in the event of a fatal error in our script. All insert and update operations were performed within the same transaction which was used to acquire the lock. This ensured that all changes could be rolled back if single input records were invalid. The input data were prepared for insert and update operations using functions in the Sga.pm module, which were mostly specific to single relations, but generally fulfilled three tasks. First, the input data were filtered to ensure that no duplicates were present within the inputs. Second, the input data were inserted or updated in batches to speed up calculations over the network. Third, values for foreign keys were collected for any record in the input data (also for records that were not inserted or updated) and passed on to subsequent functions. The insertSample function in the SGA.pm module had another important feature: the “number of run and barcode” in the measurement file was expected to reference non-control samples (Supplementary Table S1; Supplementary Methods 3.3). In order to link each sample to the water control of the respective run, insertSample internally duplicated each sample while modifying the value for the database column “iscontrol” (Supplementary Methods 5.1.1). Control samples referenced barcode 99 of the respective run (Supplementary Methods 5.3.1), as defined by the function insertSequence in SGA.pm.

Internally, importSGA.pl used the unique constraints of the relations (Supplementary Methods 5.1.1) together with the “on conflict” clause of the “insert” statement (https://www.postgresql.org/docs/14/sql-insert.html; accessed 2024/02/01) to decide if the input data needed to be synchronized with the database. In case the database was not up-to-date, the constraints determined if the input data needed to be used to create new records or update existing ones. If the only differences between the database contents and the input data were limited to columns outside of unique constraints, database records were updated. Inserts were necessary if there were differences in the columns with unique constraints. This implied that values from columns with unique constraints could never be updated with importSGA.pl (e.g. to correct a typing error), as this would have lead to the insert of an all new record. Thus, these updates had to be done manually. If database records were modified, the last action of importSGA.pl was to refresh the materialized views and to commit the changes.

In our project, FASTQ “pass” files and MetaG results were continuously added, but contents of individual files never changed. Therefore, the distinction between updates and inserts was most relevant for the measurement spreadsheet (Supplementary Methods 5.3.1) and we will use this file as an example to briefly discuss the peculiarities of automated inserts and updates with importSGA.pl. New records in the measurement file can be synchronized with the database by either supplying the full measurement file with both new and up-to-date records or a minimal measurement file containing only the new records. The latter reduces the processing time. At any rate, the measurement file needs to be formatted and columns need to be named as described earlier (Supplementary Methods 5.3.1; Supplementary Table S1). Especially all columns need to be present in the correct order, even if not all of them are needed for the insert or update. When a new patient is inserted, metadata for the patient together with his or her first sample are necessary (“patient ID,” “hospital code,” “birth date,” and “sample collection date”) (Supplementary Table S1). Patients and samples can only be added to the database, but never updated, since the aforementioned columns are part of unique constraints (Supplementary Methods 5.1.1).

The sample for a patient which is first inserted has to be the sample with the earliest sample collection date for the respective patient. Due to the static measurements which are only connected to the very first sample of each patient (Supplementary Methods 5.1.1; Supplementary Table S3), it is not possible to add an earlier sample later on. Measurements with empty values are not stored in the database. Thus, a minimal sample has no entries in the *measurement* relation, but the expected measurement names are already inserted into the *type* relation (Supplementary Methods 5.1.1). Measurements can be inserted by providing non-empty values in the input file together with the respective patient and sample metadata. Notably, the values of some columns are translated by the insertMeasurement function of SGA.pm before insertion into the database (columns with no shading in Supplementary Table S1 whose descriptions contain bullet points). These columns are not allowed to contain any unexpected values. Otherwise, values can be freely changed to update the respective *measurement* records, but changes to values of static measurements (Supplementary Table S3) are limited to the first sample. Once inserted, measurements cannot be deleted by emptying their value in the input file. Values for any columns representing derived measurements (Supplementary Table S4) are ignored, as these are calculated in the database and not read directly from the input (Supplementary Methods 5.1.1 and 5.1.2; Supplementary Table S7). The same applies to the “time point” column (Supplementary Table S7).

It is mandatory to provide the directory with FASTQ “pass” files and MetaG results to importSGA.pl. Nevertheless, if the column “number of run and barcode” (Supplementary Table S1) does not contain any (matching) patterns, no sequences and classifications are inserted. Sequences are added without classifications, if the latter is not available. However, if classifications are supposed to be added to existing sequences in the database, sequences and analysis (columns: “program” and “database,” Supplementary Table S1), patient, and sample metadata need to be provided again. Once sequences are added, the value in “number of run and barcode” (Supplementary Table S1) for the respective samples should remain unchanged: sequences can only be added to the database, even if an empty, different, or non-matching pattern is provided in “number of run and barcode.” In general, values in this column should only be changed in order to reference the original FASTQs in a different location. In this case, importSGA.pl recognizes and skips the duplicates. Similar considerations apply to the MetaG classification files. However, MetagenomicsDB supports the classification of a single read with multiple analysis programs and/or databases (Supplementary Methods 5.1.1). Thus, the value in “number of run and barcode” can be safely changed to reference new classifications, provided that the same sequences were referenced and the “program” and/or “database” columns (Supplementary Table S1) are changed simultaneously. This implies that “program” and “database” cannot be updated to correct typing errors as explained earlier for patients and samples.

#### 5.4 Portability assessment

In the following, we briefly describe which features of MetagenomicsDB can be directly applied out-of-the-box for similar studies and which need adaptation. This list is not conclusive, but should rather provide a general impression. MetagenomicsDB was designed to serve as a template for microbiome studies with a similar setup. Thus, the import and export units (Figure 1) are modular and tests are used to verify the expected behavior of modules (commit: 68b05c6; https://github.com/ IOB-Muenster/MetagenomicsDB/tree/main/bin/tests; accessed 2024/06/07). We expect that the following modules (commit: 68b05c6; https://github.com/IOB-Muenster/ MetagenomicsDB/tree/main/lib/perl/MetagDB; accessed 2024/06/07) are applicable out-of-the box for similar projects: Fastq.pm which parses the FASTQ files, Taxa.pm which parses the MetaG classifications, Table.pm which processes the measurement table, and Helpers.pm which fulfills miscellaneous functions. The same applies to Db.pm which directly interacts with the database, provided the database service name is identical (Supplementary Methods 5.3.2) and our aforementioned schema (Supplementary Methods 5.1) is used. A few adjustments have to be made to the export module Export.pm (commit: 68b05c6; https://github.com/IOB-Muenster/MetagenomicsDB/blob/main/lib/perl/MetagDB/ Export.pm; accessed: 2024/06/07) and the associated exportSGA.pl script (Supplementary Methods 5.2), depending on the available metadata, taxa, and user preferences (Section 3.3). The export to MicrobiomeAnalyst also requires the module Sga.pm which bundles project-specific functions that rely on our database schema and input data. Thus, it is not expected to be directly applicable to different projects. For example, the getMeta function in the module determines which metadata are exported to MicrobiomeAnalyst (Supplementary Table S5), the parseTable function contains the columns which are supposed to be extracted from the measurement table (Supplementary Table S7), and the import and update routines of the stored study data are heavily dependent on the database schema (Supplementary Methods 5.3.2). Mainly due to its dependency on Sga.pm, our main program that synchronizes the input data with the database, importSGA.pl, also has to be considered project-specific.

By using a relational database management system, we aim to keep the user interface inert to alterations of the physical and logical structure of the stored data (Codd 1985, ID/8). Furthermore, PostgreSQL supports the ACID principles, which ensure the integrity of the stored data (Section 4). Additionally, we took care to design a flexible database schema. For instance, any number of measurements for a sample are supported by the schema and measurement names are not predefined in the schema, but normalized via the *type* relation (Supplementary Methods 5.1.1). However, expected names are given in Sga.pm and some measurements have predefined values (Supplementary Methods 5.3.2). We expect that most relations can be reused, provided that the project is sufficiently similar. Nevertheless, the views are necessarily more project-specific: a later project might not require the calculation of weight-for-age z-scores or BMIs. Our web interface (https://www.bioinformatics.uni-muenster.de/tools/metagenomicsDB; accessed 2024/01/22) allows us to operate all units of MetagenomicsDB (Figure 1) and uses custom JavaScript and Cascading Style Sheets. Thus, the interface itself and the associated code that interacted with the database are specific to our computing environment and are not designed to be portable.

